# Inhibition of Notch signaling rescues cardiovascular development in Kabuki Syndrome

**DOI:** 10.1101/489757

**Authors:** Maria de los Angeles Serrano, Bradley L. Demarest, Tarlynn Tone-Pah-Hote, Martin Tristani-Firouzi, H. Joseph Yost

## Abstract

Kabuki Syndrome patients have a spectrum of congenital disorders, including congenital heart defects, the primary determinant of mortality. Seventy percent of Kabuki Syndrome patients have mutations in the histone methyl-transferase *KMT2D*. However, the underlying mechanisms that drive these congenital disorders are unknown. Here, we generated and characterized a zebrafish *kmt2d* null mutant that recapitulates the cardinal phenotypic features of Kabuki Syndrome, including microcephaly, palate defects, abnormal ear development and cardiac defects. The cardiovascular defects consist of abnormal aortic arches and hypoplastic ventricle, driven by previously unknown aberrant endocardial and endothelial vasculogenesis. We identify a regulatory link between the Notch pathway and Kmt2d during vasculogenesis and show that pharmacological inhibition of Notch signaling rescues the cardiovascular phenotype in zebrafish Kabuki Syndrome. Taken together these findings demonstrate that Kmt2d regulates vasculogenesis, provide evidence for interactions between Kmt2d and Notch signaling in Kabuki Syndrome, and suggest future directions for clinical research.

## Introduction

Kabuki Syndrome type I (KS, OMIM#147920) is a rare multi-systemic disorder, manifested by craniofacial anomalies including cleft lip/palate and microcephaly, hearing loss, neurodevelopmental defects, epilepsy, skeletal and skin abnormalities, and congenital heart defects (CHD) (1) (2–4) (5–7) (8) (9–11). While there is variable expressivity of the clinical hallmarks (8,12), CHD is present in ∼70% of KS patients, with a unique predilection for left-sided obstructive lesions, including hypoplastic aortic arch, coarctation of the aorta and hypoplastic left heart syndrome (13–17). *De-novo* pathogenic variants in Histone-lysine N-methyltransferase 2D (KMT2D) are causative in up to 76% of KS patients (18,19). KMT2D, also known as MLL4 and MLL2 in humans and Mll2 in mice, belongs to a family of histone 3 lysine 4 methyltransferases (20), encodes a large protein with multiple domains, and plays critical roles regulating gene expression through epigenetics mechanisms. Despite great progress in genetic diagnosis for KS, our understanding of KS phenotypic variability and the downstream molecular pathways underlying the abnormal development of specific organ systems in KS is limited. Thus, characterizing the molecular mechanisms that drive phenotypic variation of KMT2D-dependent diseases is crucial for designing therapies to ameliorate these disorders.

Germline knockout of *Kmt2d* in mice is embryonic lethal, limiting the ability to model and study KS (21). Conditional *Kmt2d* knockout in murine cardiac precursors and cardiomyocytes indicated that KMT2D is essential for regulating cardiac gene expression during heart development, primarily via di-methylation marks in lysine 4 of histone 3 (H3K4) (22). The observation that cardiac progenitor-specific *Kmt2d* deletion mutants manifest more severe forms of CHD compared to myocardium-specific *Kmt2d* deletion suggests an unexplored critical role for Kmt2d in endocardium or endothelial lineages. Understanding the contribution of KMT2D to endocardial and endothelial development is crucial given that the prognosis of KS patients depends on the diagnosis and management of left-sided obstructive cardiovascular lesions (23).

Morpholino knockdown of Kmt2d in zebrafish revealed gross neurological defects and anomalies in cardiac looping, suggesting that zebrafish might serve as a model for KS (24). However, these phenotypes are common in morpholino treatments (25–27). A recent study revealed a link between RAS/MAPK pathway hyperactivation and the neurological and craniofacial defects in the context of KS knockdown in zebrafish (28). In line with this, chemical inhibition of a downstream target of this pathway (BRAF inhibitor), partially rescued the craniofacial and neuroanatomical phenotype of *kmt2d*-depleted zebrafish larvae in transient knockdown and *kmt2d*^+/-^ heterozygous crosses (29). These findings suggest a pathway involved in some aspects of KS-neurological defects and establishes the utility of zebrafish for drugs screening in KS. However, modeling cardiovascular developmental defects in KS and their molecular pathways, and possible approaches to reduce cardiovascular defects, the major cause of death in KS, have not been explored.

Although the molecular signatures driving KS phenotype have been reported (21,24,30), it remains unclear how KMT2D impacts cardiovascular patterning. Here, we generated a zebrafish germline genetic mutant for *kmt2d* and validated it as model for KS by analyzing multiple cardinal clinical manifestations of this syndrome, including variable expressivity, short body length, palate defects, abnormal ear development and heart defects. We identify for the first time a critical role for Kmt2d in vasculogenesis and a regulatory link between Kmt2d and Notch signaling. Moreover, pharmacological inhibition of notch signaling rescues the cardiovascular defects observed in *kmt2d* mutant zebrafish, providing a platform for small molecule therapies to ameliorate the cardiovascular defects observed in KS patients.

## Results

### Zebrafish *kmt2d* null mutants exhibit phenotypes observed in human Kabuki Syndrome patients

Zebrafish *kmt2d* (ENSDARG00000037060) on chromosome 23 contains 53 exons, with a 17.6 kb mRNA encoding a 362.7 KDa protein. Although the zebrafish Kmt2d protein (UniProt E7F2F7) only has 44.3% amino acid identity with the human KMT2D (UniProt O14686), BLAST analysis of the individual protein domains showed that the Plant Homeo Domain (PHD) located at the N-terminus of the zebrafish Kmt2d has 86.1% identity with the human PHD (UniProt O14686: aa 5030 to 5137 and 126 to 217) and the SET and Post-SET domains located at the C-terminus of the zebrafish protein have 99.1% and 100% identity with the human SET (O14686: aa 5398 to 5513) and Post Set domains (O14686: aa 5521 to 5537), respectively. Important for functional analysis using reverse genetic approaches, this is the only ortholog for the human *Kmt2d* gene found in zebrafish, and there are no known paralogues.

To evaluate Kmt2d functional roles in zebrafish, we used CRISPR/Cas9 genome editing to generate zebrafish *kmt2d* null mutants. Exon 8 (ENSDARE00001117370), which contains the coding sequence for a PHD domain of the Kmt2d protein (Fig. 1A), was targeted with a single guide RNA (sgRNA). On-target mutagenesis of injected embryos (F0) was confirmed by High Resolution Melt Analysis (HRMA) of the predicted target region for the sgRNA. Germline transmission of mutant alleles was confirmed by F1 genotyping. Genotyping of individual F1 adults revealed multiple *kmt2d* alleles, three of which were selected for propagation and additional analysis: *kmt2d*^*zy58*^ (Fig. 1Bbeh 1bp deletion), *kmt2d*^*zy59*^ (Fig. 1Badg, 19bp deletion) and *kmt2d*^*zy60*^ (Fig. 1Bcfi, 2bp deletion). These mutant alleles are predicted to result in premature stop codon leading to truncation in one of the PHD domain located at the N-terminus of the protein. Loss of full length Kmt2d protein was confirmed by immunohistochemistry (Supp. Fig. 1A and 1B).

**Figure 1.**
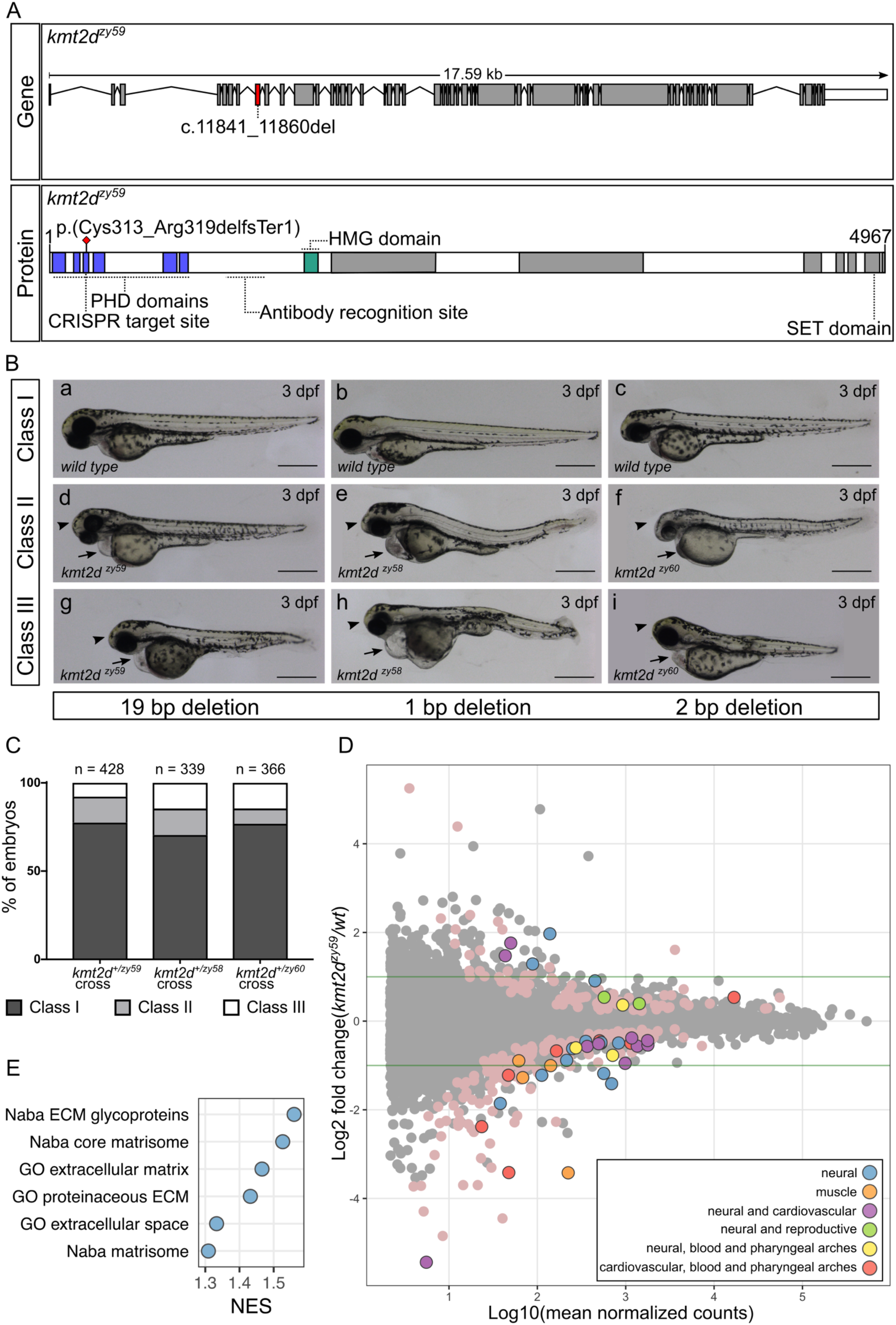
Generation and transcriptome profiling of *kmt2d* zebrafish mutants. **A.** Schematic of zebrafish *kmt2d* gene showing the 19bp deletion allele (*zy59*) and its predicted amino acid change in the protein. The gRNA was designed to target exon 8 (red shaded exon) at the 5’ end of the gene. The resulting 19bp deletion is predicted to cause an early stop codon at the level of the PHD tandem domains (blue) located at the N-terminus of the protein. The anti-Kmt2d antibody epitope is located after the PHD domains and before the HMG domain (green) allowing the validation of the early stop in *kmt2d*^*zy59*^ zebrafish null mutant. **B.** Lateral views of zebrafish *wild type* sibling embryos (A-C) and *kmt2d* mutants for 3 different alleles: *kmt2d*^*zy59*^ (A, D, G), *kmt2d*^*zy58*^ (B, E, H) and *kmt2d*^*zy60*^ (C, F, I) at 3dpf. All three alleles share the same phenotypic characteristics: microcephaly (arrowhead), heart edema (arrow) and mild to moderate body axis defects. Variable expressivity was observed in all the analyzed mutant alleles (Classes I to III). Scale bar = 500µm **C.** Variable expressivity was analyzed in three different embryo clutches resulting from a heterozygous by heterozygous cross for each mutant allele. Embryos were ranked in three different classes based on the severity of the phenotype. Class I, *wild type* and heterozygous siblings with no phenotype; class II, mutants with microcephaly and heart edema; class III, mutants with microcephaly, heart edema and shorten body axe. Percentages of different clutches were calculated per the total number of living embryos for each genotype. Chi-square test (*p = 0*.*14*) and Binomial test (*p = 0*.*09*) were performed to assess Mendelian ratios considering heterozygous embryos within the Class I category. There was no significant discrepancy between obtained and expected percentages of embryo phenotype. **D.** MA plot of differentially expressed genes from RNA-seq of individual *kmt2d*^*zy59*^ mutants (n=6) vs. *wild type* (n=6) sibling embryos at 1dpf. The log_10_ of mean normalized counts are plotted against the log_2_ fold changes for each gene tested. Green horizontal lines represent 2-fold change differences. Negative log_2_ fold changes represent genes with reduced expression in the mutants relative to wild type. Both light-pink points (no outline) and color-coded points (with outline) represent significantly differentially expressed genes (FDR adjusted p-value < 0.05). The top 50 genes (ranked by FDR adjusted p-value) were classified into 6 categories based on expression data, bibliography, phenotype information and gene ontology (Supplementary table 1). **E.** Gene set enrichment analysis. Analysis was performed by converting zebrafish gene names to human gene names using genes with a one-to-one ortholog relationship retrieved from Ensembl Compara database. The number of resulting genes identifiers analyzed was 9128 out of 33737 (Genome build Zv9, Ensembl annotation released version 79). Adjusted p-values were calculated per category. Normalized Enrichment Score (NES) of gene sets with a FDR of 5% are plotted to summarized GSEA results. PHD: Plant Homeo Domain; HMG: High Mobility Group domain; SET: Su-Enhancer-of-zeste and Trithorax domain (Histone methyltransferase activity); CNS: Central Nervous System

In order to assess the phenotypes of *kmt2d* mutations and to determine whether any identified phenotypic traits followed expected Mendelian ratios, we performed 3 different het by het crosses per allele and screened for phenotypes, with an emphasis on organs and systems that are affected in KS patients. Interestingly, embryonic development was grossly normal until 2 days post fertilization (dpf). Starting at 3dpf, embryos showed signs of cardiac edema (Fig. 1B, arrow), microcephaly (Fig. 1B, arrowhead) and body axis defects (Fig. 1Bghi), attributes similar to KS clinical manifestations. Whole body edema was evident at 3-4 dpf (Supp. Fig. 2ABC), and these phenotypes continued to develop until approximately 7 dpf (data not shown).

**Figure 2.**
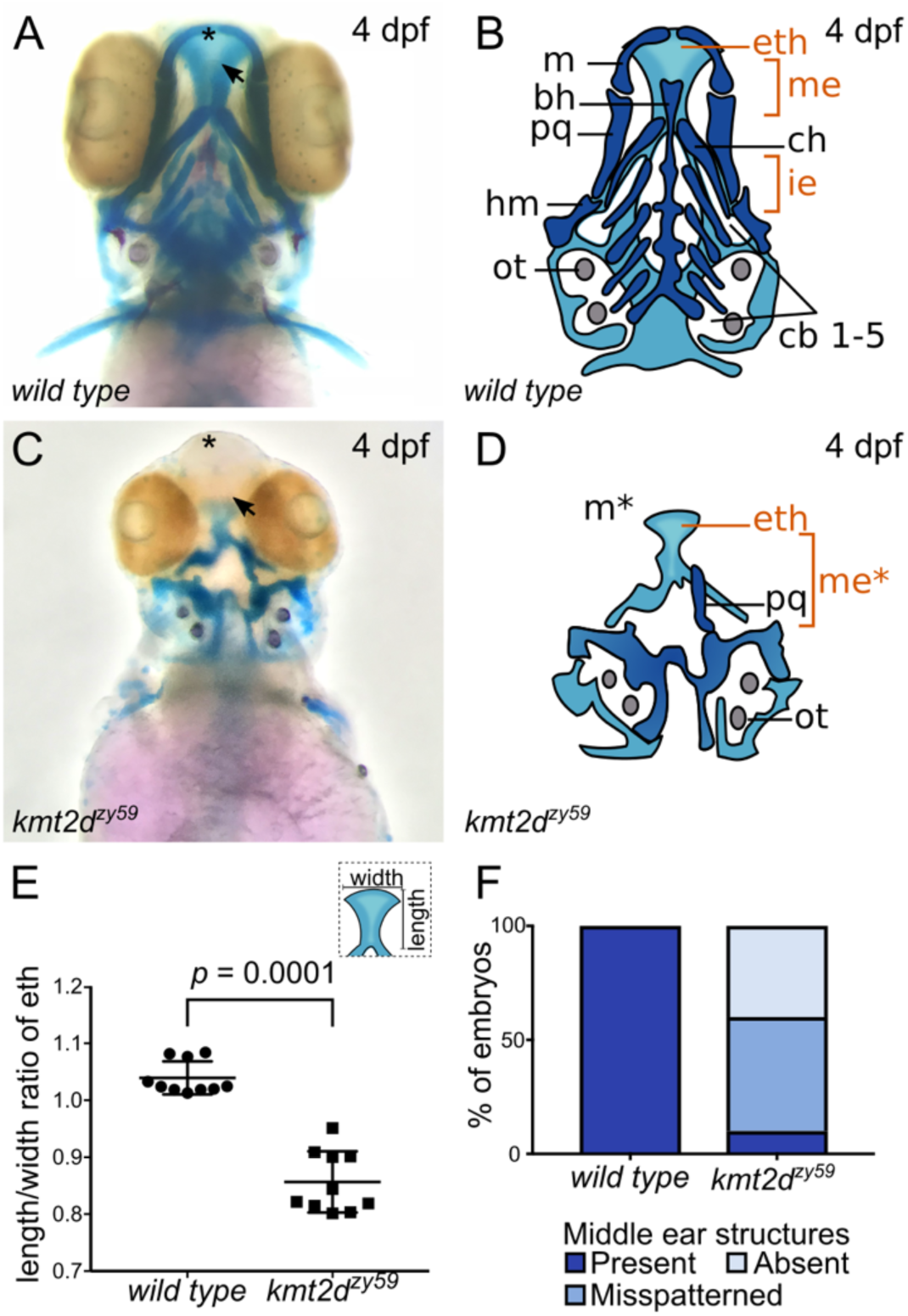
*kmt2d* mutants exhibit anomalous palate development and middle ear structures defects. **A-D.** Alcian Blue/Alizarin Red staining of cartilage and bone showing zebrafish homologous structures for mammalian palate (neurocranium) and middle ear (jaw structures). Ventral view of zebrafish sibling control (A) and *kmt2d*^*zy59*^ mutant (C) embryos at 4 dpf with corresponding simplified cartoons (B, D). *kmt2d*^*zy59*^ mutants show severe hypoplasia of visceral cartilages (C; dark blue, D) and neurocranium (C; light blue, D) when compared with *wild type* sibling (A, B). The ethmoid plate is present but displays abnormal development (eth). The cartilages that pattern the jaw in the mandibular (m, pq) and hyoid (ch, hm) arches are absent or drastically reduced. Pharyngeal arches are absent (cb1-5). The specific structures that are considered mammalian homologs of palate and middle ear are highlighted in orange (eth, me). **E.** Quantification of with/length ratio of the ethmoid plate in *wild type* siblings and *kmt2d*^*zy59*^ mutants. In all mutant embryos analyzed, the ethmoid plate was present but with a significantly reduced length/with ratio when compared with its *wild type* siblings. Statistical analysis was carried out using two-tailed t-test, *p* < 0.0001, n = 10 per genotype. **F.** Embryos were classified according to the degree of development of homolog structures for mammalian middle ear. The categories were: Present, mispatterned or absent. Qualitative assessment was plotted for percentage of embryos per genotype (n= 10 per genotype). m, Meckel’s; bh, basihyal; pq, palatoquadrate; hm, hyomandibula; ot, otolith; cb, ceratobranchial; ch, ceratohial; eth: ethmoid plate; me, middle ear mammalian homologs.

To assess whether these phenotypes segregated with the mutant alleles, we selected 16 embryos with wild-type phenotypes and 32 embryos with KS phenotypes at 3dpf and processed them individually for DNA extraction and HRMA analysis. All embryos with KS phenotypes were homozygous for the *kmt2d* mutant alleles, while embryos that appeared phenotypically normal were homozygous *wild type* or heterozygous at the *kmt2d* locus. Moreover, the percentage of embryos from a het by het cross with an abnormal phenotype was approximately 25% in 3 different clutches for each of the three alleles assayed (Fig. 1C). These results were verified by performing Chi-square and Binomial distribution tests against an expected Mendelian ratio of 75% *WT* and heterozygous embryos (no phenotype) and 25% mutant embryos (with phenotype) (Fig. 1C).

Humans with KS have wide variation in phenotypes (7,31). Similarly, zebrafish *kmt2d* mutants had a range of variable expressivity of phenotypic traits. To assess whether the observed variable expressivity occurred for different parental crosses and mutant alleles, we defined three phenotypic categories as follow: class I, WT phenotype corresponding to homozygous *wild type* and heterozygous siblings (Fig. 1Babc); class II, homozygous mutant embryos exhibiting heart edema and microcephaly (Fig. 1Bdef); class III, homozygous mutant embryos exhibiting heart edema, microcephaly and shorter body axis (Fig. 1Bghi). Embryos from 3 different parental crosses for each of three mutant alleles were screened for each category. Our results showed that, variable expressivity of the phenotypic traits is a consistent characteristic in zebrafish *kmt2d* homozygous mutants (Fig. 1C).

Our protein expression data showed that Kmt2d is strongly and ubiquitously expressed during development (Supp. Fig. 1C-F), consistent with phenotypes in multiple organ systems. In order to discover pathway perturbations that lead to KS phenotypes, we pursued the *kmt2d*-dependent alterations in transcriptional profiles that precede the onset of the KS phenotypes. RNA-seq analysis was performed on single 1 dpf embryos, prior to the appearance of a KS phenotype. The transcriptome was analyzed in 6 individual homozygous *kmt2d*^*zy59*^ mutants and 6 individual homozygous *wild type* siblings (total n = 12). For gene differential expression analysis we performed DESeq2 using Likelihood Ratio Test (LRT) (32) at a FDR of 5%. Our results showed that at 1dpf 276 genes were differentially expressed in *kmt2d*^*zy59*^ mutants when compared with their *wild type* siblings (Fig. 1D, MA plot pink and colored dots). Text-mining analysis revealed that 72.1% of the top 50 genes are associated with neural and/or cardiovascular system while the remaining genes have been associated with reproductive system, muscle and pharyngeal arches development (Fig. 1D, MA plot color-coded dots; Supp. Table 1). Furthermore, gene set enrichment analysis (GSEA) revealed a small group of gene sets that were enriched in *kmt2d*^*zy59*^ mutants (Supp. Table 2). Interestingly, all the gene sets with a 5% FDR were exclusively associated with structural extracellular matrix (ECM) glycoproteins (Fig. 1E; Supp. Table 2) suggesting essential changes in ECM composition or topography even before phenotype manifestation.

Consistent with our gross examination of *kmt2d* mutants at 3 dpf, transcriptome analysis supports the hypothesis of a relatively small group of Kmt2d-dependent genes affected early in development that could explain the molecular etiology of *kmt2d*^*zy59*^ phenotype observed at later stages. Together, these data introduce a zebrafish *kmt2d* null mutant model and demonstrate its phenotypic and molecular utility as a model for studying Kabuki Syndrome.

### *kmt2d*^*zy59*^ mutants exhibit anomalous palate development and middle ear structural defects

Our morphological and transcriptome analyses suggested that abnormal pharyngeal arch development might contribute to the spectrum of cranial defects observed in *kmt2d*^*zy59*^ embryos (Fig. 1BD). Additionally, palate and middle ear structures are derived from pharyngeal arches and are affected in KS patients, serving as diagnostic features for the syndrome (2,4–6). Zebrafish anterior neurocranium and lower jaw structures are well established models for mammal palate and middle ear development, respectively (33). To assess whether Kmt2d loss affects pharyngeal arch development, we analyzed craniofacial skeleton phenotypes by Alcian Blue/Red Alizarin staining in *kmt2d*^*zy59*^ mutants and *wild type* siblings. Craniofacial cartilage architecture is strongly affected in *kmt2d*^*zy59*^ mutants (Fig. 2) and in *kmt2d*^*zy58*^ and *kmt2d*^*zy60*^ mutants (Supp. Fig. 2DEF).

The neurocranium (Fig.2 B, D; light blue) is underdeveloped and the ethmoid plate (henceforth palate) failed to grow distally (Fig. 2AC, arrow). For *wild type* siblings, the average length/width ratio of the palate was 1.03 (Fig. 2E; n=10) while *kmt2d*^*zy59*^ mutants exhibited a significantly shorter and wider configuration with an average length/width ratio of 0.85 (Fig. 2E; n=10). These results demonstrate that Kmt2d is essential for normal palate development in zebrafish embryos. Comparably, visceral cartilages were significantly affected in *kmt2d*^*zy59*^ embryos (Fig. 2BD; dark blue). The pharyngeal arch derived structures that contribute to the support of the delicate gill tissues (Fig. 2B-D; cb1-5) are completely absent in *kmt2d* mutants. The cartilage that patterns the lower jaw was absent (Fig. 2, C, D; Meckel’s cartilage, asterisk) or drastically reduced (Fig. 2CD; palatoquadrate, hyomandibula, ceratohyal). This represents mispatterning of the structures that are homologous to the mammalian middle ear (Fig. 2D; marked as me* in orange). To evaluate reproducibility of this phenotype, we categorized embryos based on the degree of development of Meckel’s, palatoquadrate and hyomandibula cartilages and calculated percentage of individuals in each category in *wild type* siblings and *kmt2d*^*zy59*^ embryos (Fig. 2F). Our results showed that 90% of *kmt2d*^*zy59*^ embryos had absent or misspatterned cartilages in the lower jaw. Together, these results demonstrate that Kmt2d is required for normal development of palate and lower jaw in zebrafish, consistent with clinical findings in KS patients.

### *kmt2d*^*zy59*^ mutants have hypoplastic hearts and occluded lumens due to aberrant endocardial cell morphology

Congenital heart disease (CHD) is diagnosed in 28% to 80% of Kabuki Syndrome patients (34). The spectrum of CHD is wide, with prevalence of aortic coarctation, hypoplastic left heart and other left-sided obstructive defects (14,15,34). Previous studies of a mouse cardiomyocyte conditional *kmt2d* deletion demonstrated that Kmt2d is required in cardiac precursors and cardiomyocytes (22). Additionally, zebrafish *kmt2d* morphants were reported to have heart looping defects and abnormal development of atrium and/or ventricle (35). Despite the growing body of evidence that Kmt2d functions during myocardium development, it has not yet been possible to assess the effects of Kmt2d loss in all cardiac tissues in a null mutant context. To investigate the effects of Kmt2d loss during heart development, we first asked whether Kmt2d protein was expressed in both myocardial and endocardial tissues in zebrafish heart. Whole mount immunofluorescence in *wild-type* zebrafish embryos revealed that Kmt2d is ubiquitously expressed in the nucleus of both myocardium and endocardium at 2dpf (Fig. 3AB). In our *kmt2d* mutants, cardiac edema does not appear until 3 dpf, which it could be the result of an accumulative deficiency in heart morphogenesis or a secondary defect to another affected embryological process. To address whether heart edema could be the result of impaired cardiac function, heartbeat was recorded at 1, 2, 3 and 4 dpf in *kmt2d*^*zy59*^ embryos and *wild type* siblings. We found significant bradycardia in *kmt2d*^*zy59*^ embryos compared with their *wild-type* siblings (Supp. Fig. 3C; ANOVA *p* value = 0.000264, F (1,76) = 14.647). Interestingly, the mean difference between WT and mutants was constant throughout this developmental period (ANOVA, interaction effect *p* value = 0.746), suggesting bradycardia was not due to later-onset secondary effects, but is a constitutive phenotype of *kmt2d*^*zy59*^ embryos.

**Figure 3.**
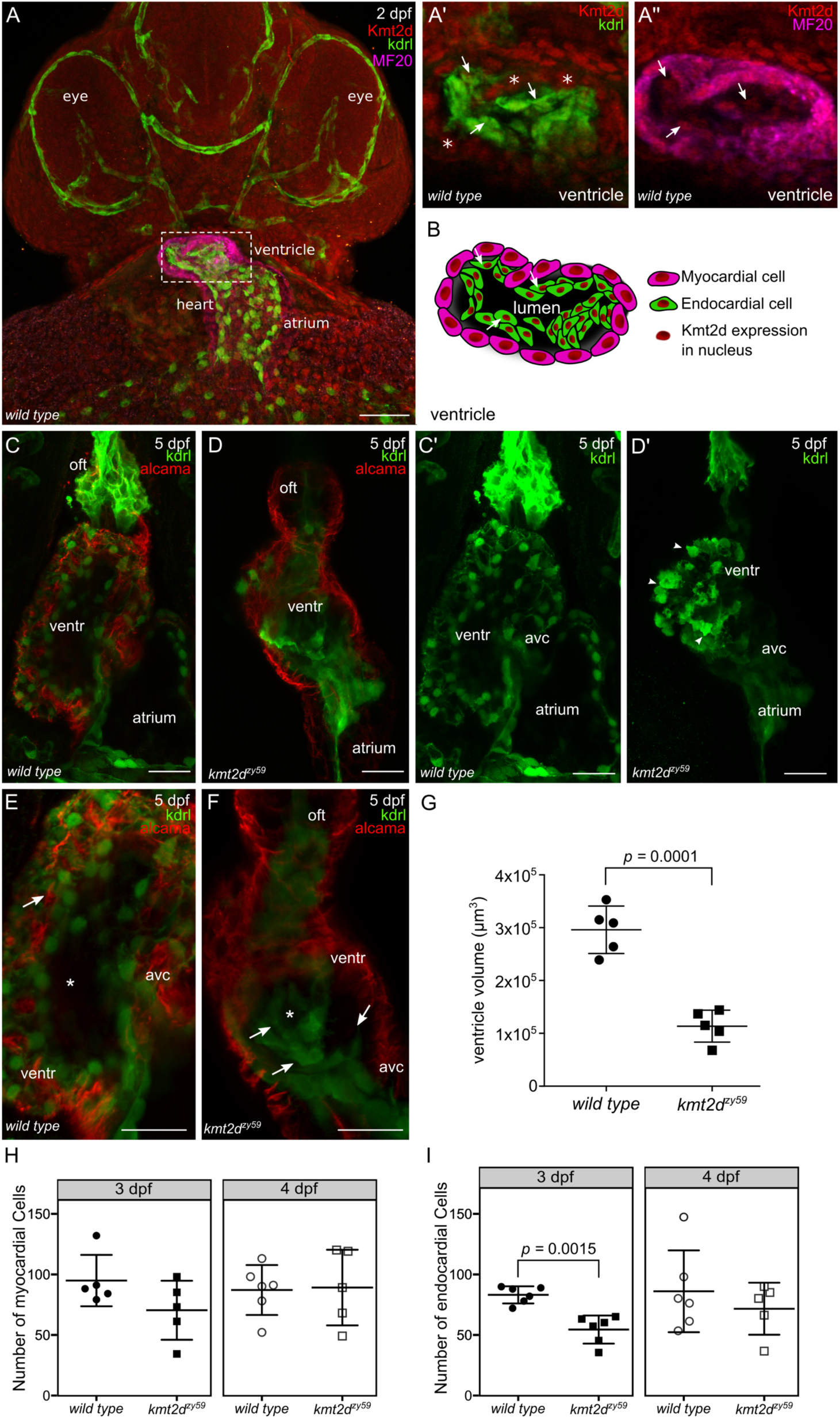
*kmt2d* mutants exhibit hypoplastic heart and aberrant endocardial cells morphology. **A.** Confocal images of whole mount immunofluorescence for *wild type* zebrafish Kmt2d protein expression at 2dpf (ventral view). Kdrl (endothelium and endocardium) and MF20 (myosin) were use as context marker for cardiovascular tissues and myocardium. Kmt2d expression is found in the nuclei of myocardial (A’, asterisk in example cells) and endocardial (A’’, arrows) cells in zebrafish heart (A’ and A” are zoomed images in ventricle area). Scale bar = 100µm **F.** Cartoon showing Kmt2d nuclear expression in both, myocardium and endocardium tissue of zebrafish heart. Arrows in A’, A’’ and F are showing Kmt2d endocardial expression in the same set of cells. **C-D.** Confocal images of wild type *tg(kdrl:GFP)* sibling (C, C’) and *kmt2d*^*zy59*^;*tg(kdrl:GFP)* (D, D’) embryos at 5dpf. Ventral view. *kmt2d*^*zy59*^ mutant show aberrant cardiac morphology with significantly reduced ventricle size (D). Myocardial cell labeled with alcama antibody (C-D, red) show normal cell morphology in both genotypes. Maximum intensity projections of GFP channel evidenced abnormal morphology of endocardial cells (D’). Some endocardial cells exhibited cell protrusions (D’, arrowheads). **E, F.** High magnification images of the heart ventricle chamber in a middle-plane view from the 3-dimentional dataset in *wild type* sibling embryo (E, higher magnification of C) and *kmt2d*^*zy59*^ mutant (F, higher magnification of D) at 5dpf. Z-stack analysis of the dataset revealed that endocardial cells from the ventricle are organized in concentric layers (F, arrows) filling up the cardiac lumen in *kmt2d*^*zy59*^;*tg(kdrl:GFP)* mutants (E, F, asterisk). **G.** Ventricle cavity volume measurements in 5 embryos per genotype. Statistical analysis was carried out using two-tailed t-test, *p* < 0.0001. References: oft, outflow tract; ventr, ventricle; avc, atrio-ventricular canal. Scale bar = 25µm. **H, I.** Ventricle myocardial (H) and endocardial (I) cells quantification at 3 and 4 dpf in zebrafish *wild type* siblings and *kmt2d*^*zy59*^ mutants. Embryos were processed for immunofluorescence against myosine heavy chain (MF20) and GFP (Kdrl). Nuclei were stained with DAPI for cell quantification with Imaris software. **H.** Myocardial cells, t-tests per time point, *p* values as follow: 3 dpf *p* value = 0.129, effect size = 24.6. 4 dpf *p* value = 0.906, effect size = -2. Bonferroni corrected p values for 2 t-tests *p adjusted* values, 3 dpf = 0.258 and 4dpf = 1.000. **I.** Endocardial cells, t-tests per time point, *p* values as follow: 3 dpf *p* value = 0.0007, effect size = 29. 4 dpf *p* value = 0.4139, effect size = 14.6. Bonferroni corrected *p* values for 2 t-tests *p adjusted* values, 3 dpf = 0.0015 (value reported in figure) and 4dpf = 0.8278.

Next, we analyzed myocardium and endocardium patterning using *tg(kdrl:GFP) crossed into the kmt2d*^*zy59*^ *line* to mark endocardium and endothelium. Cardiac morphology was significantly altered in *kmt2d*^*zy59*^ at 5dpf, with overall heart size reduced and a hypoplastic ventricle (Fig. 3CD). To assess whether the hypoplastic ventricle was due to abnormal myocardium development, we analyzed ventricle cardiomyocyte cell numbers at 3 and 4 dpf in *wild type* siblings and *kmt2d*^*zy59*^ mutants. Additionally, myocardial cell architecture was evaluated with the marker Alcama at 3 dpf. The numbers of ventricle myocardial cells in *wild type* siblings and *kmt2d*^*zy59*^ mutants were equivalent (Fig. 3H; t-tests per time point, *p* values as follow: 3 dpf *p* value = 0.129, effect size = 24.6. 4 dpf *p* value = 0.906, effect size = -2. Bonferroni corrected p values for 2 t-tests *p adjusted* values, 3 dpf = 0.258 and 4dpf = 1.000.), suggesting that the hypoplastic ventricle phenotype in zebrafish *kmt2d*^*zy59*^ is not due to a reduction in the myocardial cell population.

At 3 dpf, zebrafish heart ventricle undergoes the tightly regulated process of trabeculation (36,37), whereby myocardial cells extrude and expand into the lumen of the ventricle (38) forming a network of luminal projections consisting of myocardial cells lined by endocardial cells (38,39). After this process, myocardial cells of the outer ventricle curvature have a characteristic elongated shape (40). In order to assess whether cardiomyocyte cell shape was affected in *kmt2d*^*zy59*^ mutants, we measured outer curvature ventricle myocardial cell area and sphericity in *wild type* siblings and *kmt2d*^*zy59*^ at 3 dpf as previously described (40). Our results showed that ventricle myocardial cells morphology is equivalent in *wild type* siblings and *kmt2d*^*zy59*^ mutants at 3 dpf (Supp. Fig. 3A, t-test *p =* 0.583 n.s. t=0.59 dF=5 for area; Supp. Fig. 3D t-test, *p* = 0.946 n.s.t=0.71, dF=6 for myocardial cells sphericity). Overall these results suggest that the hypoplastic heart ventricle in *kmt2d*^*zy59*^ is not driven by a reduction in size or absolute numbers of myocardial cells.

In contrast to what we observed in myocardium, the endocardium of *kmt2d* mutant hearts displayed strikingly abnormal morphology (Fig. 3C’D’), with endocardial cells in the ventricle forming discrete aggregates in which some individual cells show protruding borders resembling mesenchyme-like morphology (Fig. 3D’; arrowheads). In *kmt2d*^*zy59*^ embryos, endocardial cells lose their tight interaction with the adjacent myocardium and form concentric layers (Fig. 3EF; arrows) that ultimately occlude the lumen of the ventricle (Fig. 3EF; asterisk). Thus, endocardium mispatterning led to lumen volume reduction in all analyzed mutant samples (Fig. 3G; t-test *p* = 0.0001 t=7.54, dF=8). Interestingly, quantification of ventricle endocardial cells in *wild type* vs. *kmt2d*^*zy59*^ mutants showed a significant reduction in the number of EC cells in *kmt2d* mutants (Fig. 3I, t-tests per time point, *p* values as follow: 3 dpf *p* value = 0.0007, effect size = 29. 4 dpf *p* value = 0.4139, effect size = 14.6. Bonferroni corrected *p* values for 2 t-tests *p adjusted* values, 3 dpf = 0.0015 and 4dpf = 0.8278). However, this reduction in endocardial cell number did not appear to be due to cell death, since anti-active-caspase3 IF did not find significant differences in the number of apoptotic cells in *kmt2d* mutant hearts in the ventricle (Supp. Fig. 3 B; arrowhead).

Altogether, these results indicate that Kmt2d is required for endocardium pattering during zebrafish cardiogenesis. Loss of *kmt2d* reduced overall heart size resulting in hypoplastic ventricle with occluded cavity. These results suggest a previously unknown role of Kmt2d in endocardial patterning might contribute the cardiac phenotype observed in KS patients.

### *kmt2d*^*zy59*^ mutants are defective in vasculogenesis and angiogenesis

While KS patients frequently manifest abnormalities of the aortic arches, including hypoplastic aortic arch or coarctation, a mechanistic understanding between *KMT2D* mutations and vascular anomalies is lacking. During early development, common progenitor cells (angioblasts) give rise to the cardiac endocardium and to the primary vascular endothelium through vasculogenesis (41–43). After the primary vascular plexus is formed, a more complex vascular network is established through angiogenesis (production of vessels from preexisting ones)(44,45). Considering their common developmental origin and our results demonstrating abnormal endocardium patterning, we assessed whether *kmt2d*^*zy59*^ mutants have normal vascular patterning. To do so, we analyzed general vasculature integrity through *o-dianisidine* staining (46) in wild type siblings and *kmt2d*^*zy59*^ embryos at 6dpf (Supp. Fig. 4A-D) and found vasculature mispatterning reflected by aggregates of red blood cells in the head and aortic arches (Supp. Fig. 4BD; white arrowheads).

**Figure 4.**
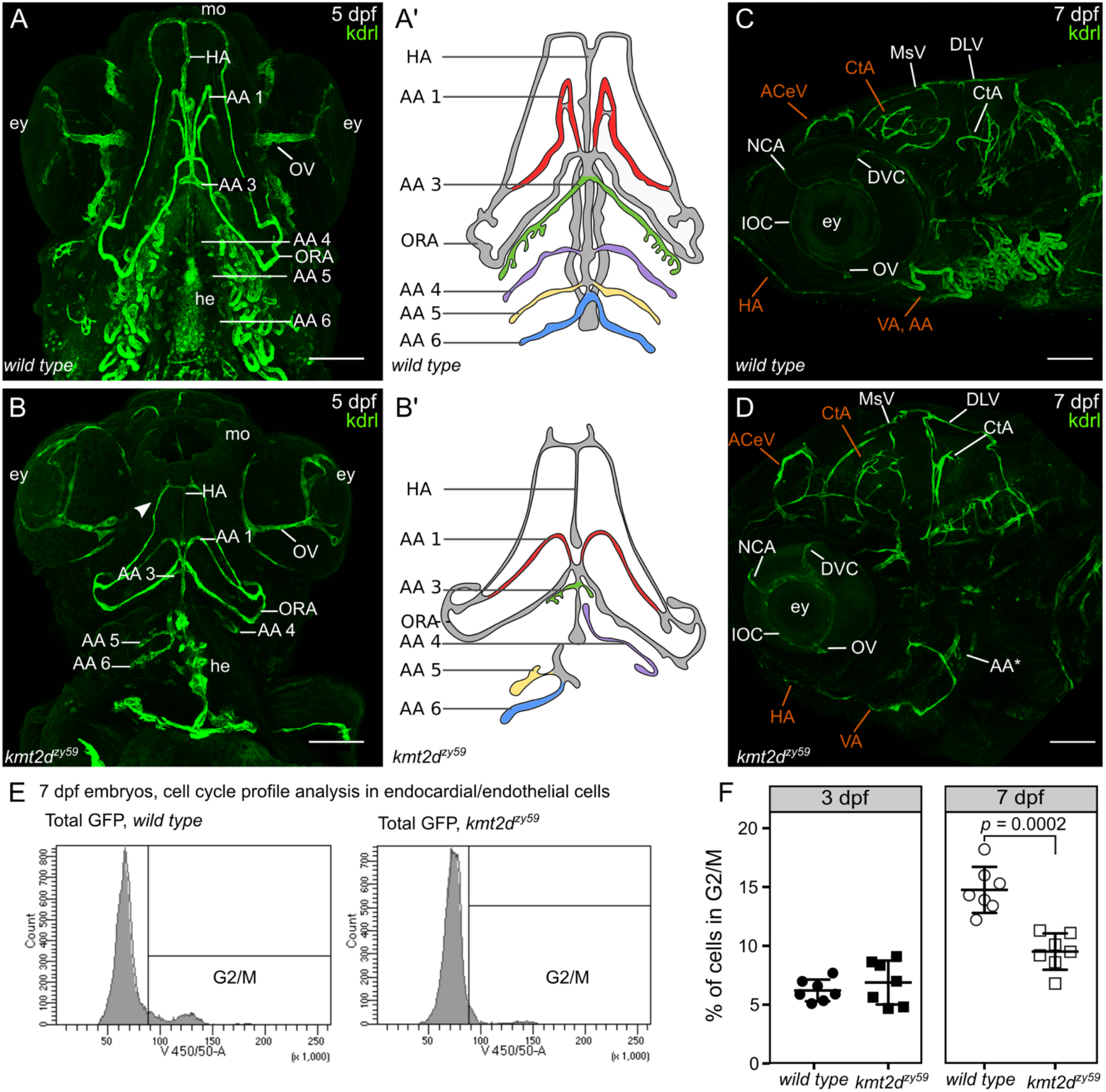
*kmt2d*^*zy59*^ mutants fail to develop aortic arches and exhibit misspatterned cephalic vasculature. **A-B.** Ventral view of vasculature in *wild type tg(kdrl:GFP)* sibling (A) and *kmt2d*^*zy59*^;*tg(kdrl:GFP)* mutants at 5 dpf. Cephalic vascular architecture in *kmt2d*^*zy59*^ embryos is abnormal with reduced elongation in the anteroposterior body axis, loss of bilateral symmetry and reduced hypobranchial artery (A, B; HA). In *kmt2d*^*zy59*^ mutants the mandibular arch (A, B; AA1) is shorter with minimum elongation towards the anterior area of the embryo, the third branchial arch (A, B; AA3) is rudimentary while the posterior branchial arches are reduced or absent (A, B; AA4-6). Mouth and eyes in *kmt2d*^*zy59*^ show primitive characteristics with differences in the thickness of the optic vessel in particular (A-B, OV). Note the abnormal endocardial component of the heart (A, B; he). A’ and B, are simplified cartoons of the main differences in the aortic arches development in *wild type* (A’) and mutant (B’) backgrounds. Branchial vasculature loops were removed to allow better visualization of aortic arch points of origin. **C-D.** Lateral view of cephalic vasculature in *wild type tg(kdrl:GFP)* sibling and *kmt2d*^*zy59*^;*tg(kdrl:GFP)* mutants at 7 dpf. *kmt2d*^*zy59*^ exhibit a complete absence of the vascular loops associated with AA3-6 with only a rudiment of the AA3 present (D, AA*). The general cranial vascular network is mispatterned with a particular strong impact in the central artery, anterior cerebral vein and ventral aorta (D; CtA, ACeV, VA in orange). In *kmt2d*^*zy59*^ mutant, most vessels show reduced lumen with the exception of the nasal ciliary artery (C, D, NCA), dorsal ciliary vein (C, F, DVC) and inner optic circle (C, F, IOC) that show thicker vessel diameter. Note the reduced HA from a lateral view (D, HA in orange) (scale bars = 100 µm). **E.** Cell cycle profile analysis by FACS for 7 individual embryos per genotype at 7dpf. The gates set up for nuclear staining (DAPI) in kdrl positive cells (endothelial and endocardial cells) are shown. **F.** Cell cycle profiles for Kdrl positive cells in *kmt2d*^*zy59*^ mutants show no significant difference in the percentage of G2/M cells at 3 dpf. In contrast, at 7 dpf there is a significant decrease in number of dividing cells in *kmt2d*^*zy59*^ mutants when compared with *wild type* siblings. Unpaired two-tailed t-test, *p* = 0.407 n.s for 3 dpf; *p* = 0.0002 for 7 dpf, n = 7 per genotype. HA, hypobranchial artery; AA1, mandibular arch; AA2, hyoid arch; AA3, first branchial arch; AA4, second branchial arch; AA5 third branchial arch; AA6, fourth branchial arch; ORA, opercular artery; MsV, mesencephalic vein; ACeV, anterior cerebral vein; VA, ventral aorta; DLV, dorsal longitudinal vein; CtA, central artery; NCA, nasal ciliary artery; DCV, dorsal ciliary vein; IOC, inner optic circle; OV, optic vein; ey, eye.

To assess whether the vascular phenotype in *kmt2d*^*zy59*^ mutants is driven by vasculogenesis or angiogenesis defects, we analyzed vascular architecture in *kmt2d*^*zy59*^;*tg(kdrl:GFP)* mutants and *tg(Kdrl:GFP) wild-type* siblings, by focusing on vasculogenesis of aortic arches (AA) and angiogenesis of the cranial vessels network at 3, 4, 5 and 7dpf. Aortic Arch patterning, thickness and development of general vasculature is dramatically affected in *kmt2d*^*zy59*^ mutants (Fig. 4AD; Supp. Fig. 4FH) when compared with their *wild type* siblings (Fig. 4AC; Supp. Fig. 4EG). Additionally, *kmt2d* mutants exhibit a primitive mouth with underdeveloped vasculature that fails to elongate anteriorly (Fig. 4B, arrowhead). Aortic arches are vestigial or completely absent in *kmt2d* mutants. Consequently, all the vessels that branch into the gills are missing (Fig. 4B; vessels on both sides of the heart). 3D volume rendering of confocal imaging confirmed that aortic arch 1 fully forms but remains in a primitive state while aortic arches 3, 4, 5 and 6 are atrophic on one side of the left-right symmetry axis and missing on its specular side (Fig. 4A’B’ and Videos 1-2). This result indicates that while the initial formation of the vascular plexus proceeds, the overall vasculogenesis process is markedly abnormal in *kmt2d*^*zy59*^ mutants.

The vascular plexus of the brain develops with all the major vessels present but with aberrant patterning, particularly evident in the central artery (CtA), anterior cephalic vein (ACeV), hypobranchial artery (HA) and ventral artery (VA) (Fig. 4CD; orange segments indicate altered patterning). These observations suggest that even when blood vessels sprout and form, they fail to establish a normal vascular patterning (Supp. Fig. 4E-H).

We next asked whether the *kmt2d*^*zy59*^ vascular defect was driven by abnormal endothelial cell proliferation, by profiling endothelial cell cycle by fluorescent activated cell sorting (FACS). In order to assess specific genotypes, single embryos were dissociated for FACS, using *kmt2d*^*zy59*^;*tg(kdrl:GFP)* embryos and. *tg(Kdrl:GFP) wild-type* siblings at 3 dpf and 7 dpf (Fig. 4EF; Supp. Fig. 4). Strikingly, our results showed no significant difference in endothelial cell cycle profile at 3 dpf (unpaired t-test, *p* = 0.4, t=0.86 dF=12 n.s.). In contrast, at 7 dpf *kmt2d*^*zy59*^;*tg(kdrl:GFP)* embryos showed a significant reduction in proliferative endothelial cells (Fig. 4F; t-test, *p* = 0.0001, t=5.56 dF=12, n = 7 per genotype). The reduction in endothelial/endocardial cell proliferation was confirmed by confocal analysis of phospho S10 Histone 3 marker at 5dpf (Suppl. Fig. 7, A, C). These data indicate that the vascular misspatterning in *kmt2d*^*zy59*^ mutants is not driven by reduced cell proliferation at early developmental stages, when the cardiovascular system is being established, but by a reduction in proliferation at a later stage. In contrast to normal proliferation at 3 dpf, there was a strong reduction in EC proliferation at 5 dpf and 7 dpf, particularly evident in the area of the vasculature that irrigates the gills (Supp. Fig. 7A; white dashed line), which are normally stablished from 5 dpf to 7dpf through active cell proliferation from vessels derived from AA 3-6 (47). Our data suggest that the difference in cell cycle proliferation at these time points is the result of the lack of AA and, consequently, the absence of blood vessels that originate from them. Overall, these results demonstrate a key role of Kmt2d in zebrafish for vascular patterning, in both vasculogenesis and angiogenesis.

### *kmt2d*^*zy59*^ mutants fail to develop AA 3 to AA 6 before gross phenotype appearance

In zebrafish, aortic arch 1 (AA 1) forms at 24 hpf, coincident with the time when circulation begins. A vestigial AA 2 also forms at this timepoint, which is immediately replaced by the opercular artery (ORA). AA 3 to 6 emerge later, between 2dpf and 2.5 dpf (47,48). As described above, *kmt2d*^*zy59*^ mutants are able to form AA 1 and the ORA but they fail to form the subsequent aortic arches. Additionally, our data indicate that endothelial cell proliferation is normal in *kmt2d*^*zy59*^ mutants at 3dpf, the timepoint at which the gross embryological phenotypes begin appear. We explored two possible mechanisms for the absence of AA3 through AA6: (1) the first defects driving the *kmt2d*^zy59^ vascular phenotype occurs sometime after 2 dpf or (2) alternatively, AA 3 to 6 develop normally and at a later stage they become atrophic and regress. To test these possibilities, we performed live time-lapse analysis of aortic arch development in *wild type tg(kdrl:GFP)* siblings and *kmt2d*^*zy59*^;*tg(kdrl:GFP)* mutants from 2 dpf to 3 dpf.

At 2 dpf, *kmt2d*^*zy59*^ mutants have normal gross morphology but time-lapse acquisition displays thinner blood vessels and abnormal AA sprouting when compared with their wild-type siblings (Fig. 5A-D). Interestingly, *kmt2d*^*zy59*^ mutants were not able to form the primary AA sprouts during the duration of data acquisition (Fig. 5B-B’’’DD’, Video 4), while *wild type* embryos succeeded in developing and extending AA 3 through 6 (Fig. 5A-A’’’CC’, Video 3). This result suggests that *kmt2d* mutants do not form AA 3 to 6 within the same time interval as their wild-type counterpart. On the other hand, the time-lapse videos show high endothelial cell activity in the area were the AA should emerge (ventral boundary of the lateral dorsal aorta, LDA), suggesting that AA could develop at a later time point. To investigate this possibility, embryos subjected to time-lapse analysis were recovered for 5 hours and processed for IF and confocal analysis. At approximately 3 dpf, *kmt2d* mutants showed primary sprouting of abnormal AA 3 to 6 that were not present at earlier stages (Fig. 5EF). This indicates that *kmt2d* mutants eventually form atretic AA around 3 dpf that will ultimately become vestigial at later time points (Fig. 4AB).

**Figure 5.**
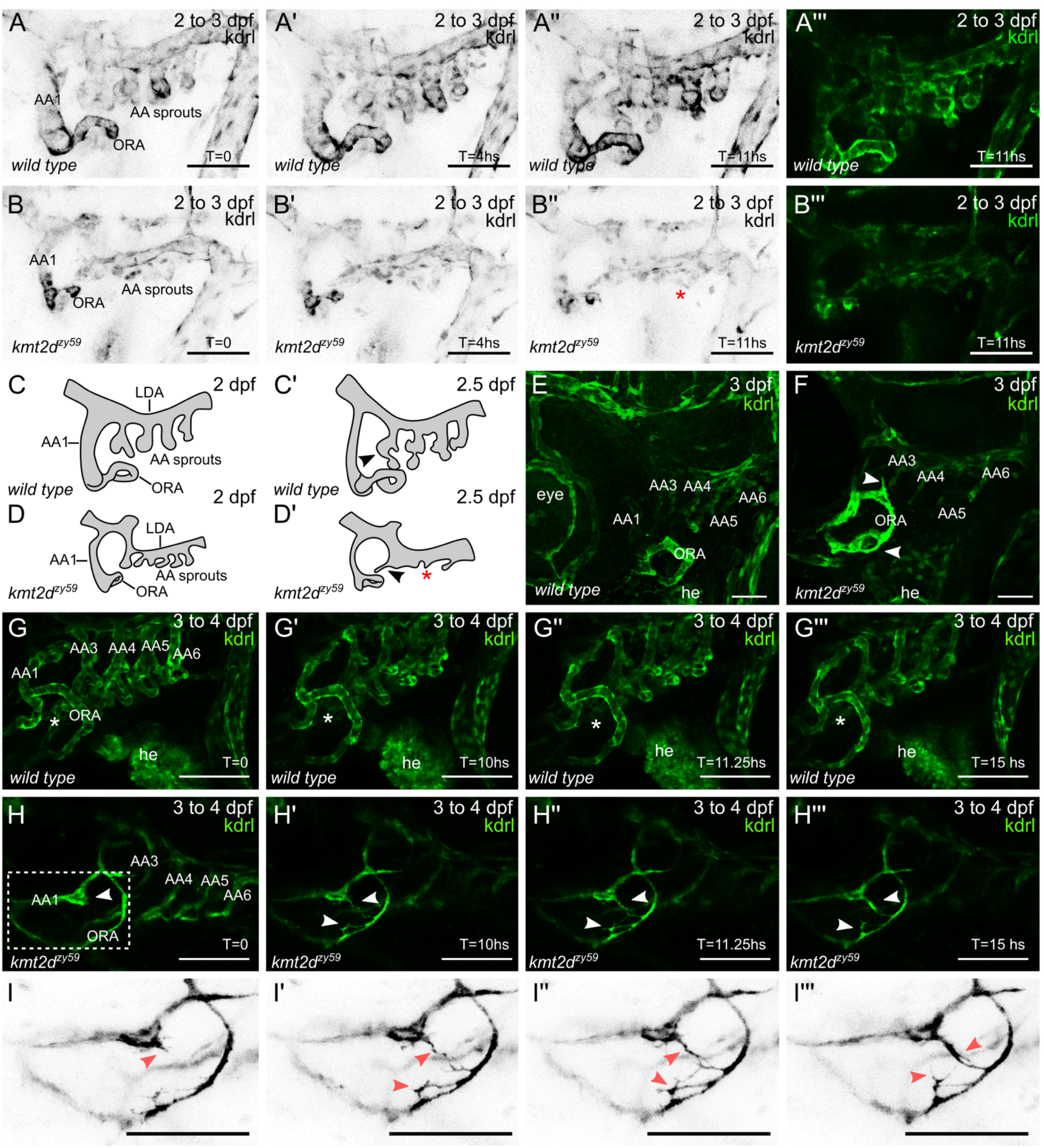
*kmt2d*^*zy59*^ mutants fail to develop AA 3 to AA 6. **A-B.** Still images (MIP) from time-lapse live imaging performed from 2 dpf to 3 dpf. Cranial-lateral view at the level of aortic arches development from *wild type* sibling (A-A’’’, Video 3) and *kmt2d*^*zy59*^ mutant (B-B’’’, Video 4). Images were selected at the 0, 4 and 11 hour time-points. Images were converted to gray scale and inverted for better visualization (A-A’’, B-B’’). A’’’ and B’’’ show A’’ and B’’ without gray scale processing. Red asterisk (B’’) denotes abnormal vascular development of AA sprouts at the level of the ventral border of LDA. (scale bars = 50 µm). **C-D**. Schematic cartoons highlighting the development of AA in *wild type* embryos (CC’) and *kmt2d*^*zy59*^ mutants (DD’). C corresponds to A, C’ correspond to A’’, D corresponds to B and D’ to B’’. Red asterisk indicates ectopic AA sprouting. **E-F**. Cranial-lateral view of vasculature in *wild type tg(kdrl:GFP)* sibling (A) and *kmt2d*^*zy59*^;*tg(kdrl:GFP)* mutants at 3 dpf. After time-lapse experiment (AB), embryos were released from agarose and processed for IF and confocal imaging. Wild-type embryos show correct patterning and secondary sprouting of AA3 to AA6 (E). At 3 dpf *kmt2d*^*zy59*^ embryos show abnormal development of primary sprouts of AA3 to AA6, which are thinner and atrophic. In contrast, AA1 and ORA are thicker and exhibit abnormal morphogenesis and endothelial cell protrusions (F, white arrowhead) (scale bars = 25 µm). **G-I.** Still images (MIP) from time-lapse live imaging performed from 3 dpf to 4 dpf. Cranial-lateral view at the level of aortic arches development from *wild type* sibling and *kmt2d*^*zy59*^ mutant (Supplementary Videos available on request). Images were selected at the 0, 10, 11.25 and 15 hour time-points. Asterisks (G-G’’’) denote normal vascular development in the area between AA1 and ORA. No blood vessel forms in this area in *wild type* background. Arrowheads (H-H’’’, I-I’’’) indicate endothelial cells extending multiple filopodia and forming a new and ectopic blood vessel. White dashed rectangle (H) specifies zoomed area in I. Images were set on gray scale and look up table was inverted for better contrast of tip cell-like endothelial cells in *kmt2d*^*zy59*^ mutant (I-I’’’) (scale bars = 100 µm). AA1, mandibular arch; AA2, hyoid arch; AA3, first branchial arch; AA4, second branchial arch; AA5 third branchial arch; AA6, fourth branchial arch; ORA, opercular artery; LDA, lateral ventral aorta; he, heart.

### Ectopic endothelial sprouting in *kmt2d*^*zy59*^ mutants

Our super resolution confocal analysis of the 3 dpf embryos found distinctive aberrant sprouting of endothelial cells from AA1 and ORA in *kmt2d*^*zy59*^ embryos (Fig. 5F, white arrowheads). This observation led us to suggests a possible angiogenesis phenotype that involves ectopic blood vessels formation in *kmt2d* mutants. To test this hypothesis we performed time-lapse analysis of aortic arch development in *wild type tg(kdrl:GFP)* siblings and *kmt2d*^*zy59*^;*tg(kdrl:GFP)* mutants at a later time point, from 3 dpf to 4 dpf. At 3 dpf (72 hpf), aberrant blood vessels phenotypes were apparent in *kmt2d* mutants, with both decreased vessel lumens and abnormal patterning (Fig. 5GH). Strikingly, by 10 hours later (82 hpf), *kmt2d*^*zy59*^;*tg(kdrl:GFP)* mutants showed ectopic endothelial cell sprouting, emerging from the AA1 and ORA (Fig. 5G G’’H’H’’; arrowheads; II’’, red arrowheads). These cells possessed many long and dynamic filopodia extending towards opposite cells with similar comportment; resembling tip cell morphology that characterizes normal angiogenesis (49). However, in the context of *kmt2d*^*zy59*^ mutants, the observed tip cell-like behavior is observed in aberrant positions, ultimately producing ectopic blood vessels (Fig. 5H’’’I’’’; Supp. videos available on request). This ectopic and abnormal endothelial cell behavior was also observed at the level of AA3 as well as in other regions of the recorded area (Supp. videos available on request), suggesting that the tip-like phenotype can potentially occur in any endothelial cell of *kmt2d*^*zy59*^ mutants.

Ectopic blood vessel formation is a well described response to hypoxia conditions (50). Considering the very limited lumen in *kmt2d* mutant blood vessels, we asked whether ectopic blood vessel formation was a consequence of hypoxia response mechanisms or if it is a direct consequence of the lack of Kmt2d affecting endothelial cell behavior. To test this, we induced hypoxia by treatment with dimethyloxalylglycineinduced (DMOG), an inhibitor of HIF-prolyl hydroxylases which thereby stabilizes Hif-1 and triggers hypoxia response genes even in normoxia conditions (51). *Tg(kdrl:GFP)* wild-type siblings and *kmt2d*^*zy59*^;*tg(kdrl:GFP)* mutant embryos were treated from 3 dpf to 4 dpf with DMOG and DMSO as control. After treatment, embryos were washed and processed for IF and confocal imaging. At 4 dpf, *wild type* embryos treated with DMOG showed ectopic vessel sprouting at the level of the optic vein (OV) (Supp. Fig. 5AB, white arrow). In contrast, AA1 and ORA were not responsive to DMOG treatment (Supp. Fig. 5AB), suggesting that these vessels have a higher resistance threshold to hypoxia. Interestingly, *kmt2d* mutant embryos treated with DMOG did not show any extra vessel sprouting (Supp. Fig. 5BD) beyond the ectopic blood vessel sprouting from the AA1 and ORA in normoxia (Supp. Fig. 5BD, white arrow), indicating that hypoxia does not increase or alter the ectopic blood vessels formed in *kmt2d* mutants. These results suggest that the aberrant endothelial cell behavior and ectopic blood vessel sprouting seen in *kmt2d* mutants are not due to hypoxia response, and that Kmt2d functions to suppress ectopic or hyperactive angiogenesis. Altogether, these results demonstrate that Kmt2d is required for the timely and normal development of AA 3 to AA6 in zebrafish.

### Notch signaling is hyperactivated in endocardial cells of *kmt2d* mutants

The tip cell-like phenotype directed us to investigate an iconic molecular mechanism involved in tip-stalk cell identity as well as in endocardium patterning during cardiogenesis, Notch signaling pathway. Multiple studies in mice, zebrafish, cell culture and tumor models have shown that Notch pathway is a key regulator of vertebrate vasculogenesis and angiogenesis (52–58). Considering our discovery of ectopic angiogenesis in *kmt2d* mutants and previous observations that Notch signaling regulates endocardial and endothelial cell growth, differentiation and patterning, we investigated whether *kmt2d* mutants have altered Notch signaling activity. We injected *kmt2d* CRISPR/Cas9 into a Notch signaling reporter line *tg(tp1:EGFP)*^*um14*^ (59) and performed F0 analysis (60) (Supp. Fig. 6) to assess Notch signaling activity in *kmt2d* mutants. At 3 dpf, Notch signaling in the zebrafish heart is strongly active in endocardial cells of the outflow tract and AV canal, with a weaker activity in some endocardial cells of the ventricle (Fig. 6A; GFP positive cells). Interestingly, in *kmt2d* mutants, the number of endocardial cells with positive Notch activity is significantly increased in both ventricle and atrium (Fig.6BC t-test, *p* = 0.0001 in ventricle t=4.95 dF=36, atrium *p* = 0.0001, t=10.99 dF= 36 n = 7 per treatment). To validate that endogenous Notch signaling is altered in *kmt2d* F0 mutants and avoid possible artifacts of transgenic reporter lines, we next tested endogenous components of the Notch pathway in *kmt2d*^*zy59*^ mutants by RT-qPCR and IF at 3 dpf. Of these components, only transcripts of Notch-specific transcription factor *rbpja* were increased in *kmt2d*^*zy59*^ mutants, while the notch1b receptor and the downstream target, *hes1*, mRNA levels were comparable to the transcripts level found in *wild type* siblings (Fig. 6D). To investigate whether elevated expression of *rbpja* mRNA resulted in elevated Rbpj protein expression, we performed immunofluorescence against Rbpj Notch transcription factor and compared protein levels in the heart in *wild type tg(Kdrl:GFP)* siblings and *kmt2*^*zy59*^*;tg(Kdrl:GFP)* mutants. At 5 dpf, Rbpj protein was present in all heart tissues with enhanced Rbpj protein accumulation in *kmt2*^*zy59*^ mutants, with well-defined nuclear localization. To quantify this elevated expression in endothelial/endocardial cells and to determine whether this increased Rbpj protein expression started at earlier stages, we performed single embryo FACS of *tg(Kdrl:GFP) wild-type* siblings and *kmt2*^*zy59*^*;tg(Kdrl:GFP)* mutants at 3dpf. Embryos were processed for immunofluorescence against GFP (driven by Kdrl in endothelial/endocardial cells) and Rbpj. After gating for single cells, samples were gated for GFP to exclusively study the endothelial/endocardial single cell population (Supp. Material 2). Subsequently we measured in EC the percentage of Rbpj positive cells and the Rbpj Median Fluorescence Intensity (MFI) as an indicative of protein expression levels (Fig. 6D). Consistent with our previous cell counting results, we observed that the overall number of Rbpj positive cells in EC from *wild type* sibling and *kmt2*^*zy59*^ mutants is equivalent at 3dpf (Fig. 5A, % of EC Rbpj positive, unpaired t-test, *p* = 0.69, t=0.42 dF=6 n.s.). Crucially, the MIF values in *kmt2*^*zy59*^ mutants were significantly higher, indicating elevated Rbpj protein levels in Kmt2-deficient endothelial/endocardial cells (Fig. 6D, t-test *p* = 0.0015, t=5.51 dF=6).

**Figure 6.**
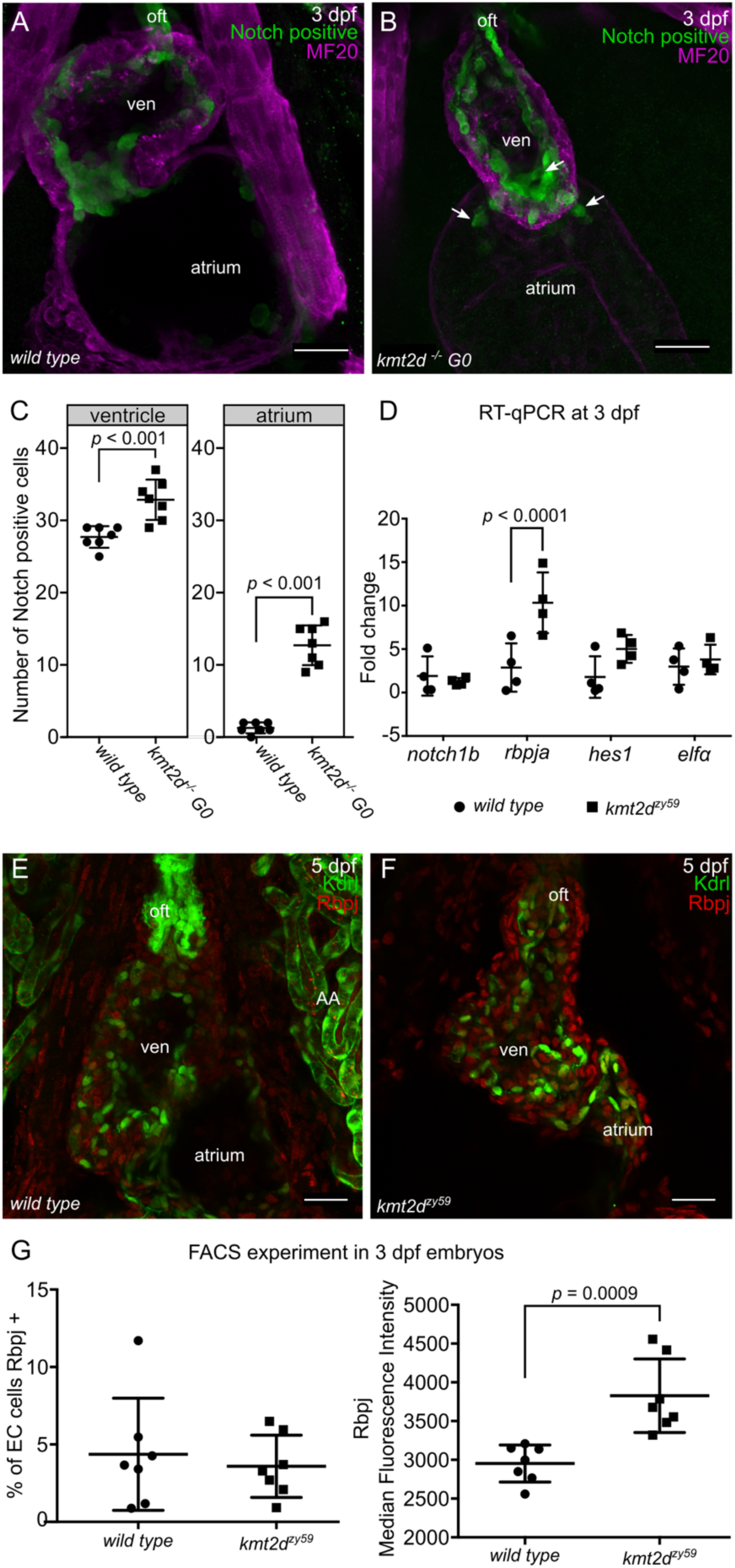
Notch signaling is hyperactive in *kmt2d* mutant endothelium/endocardium. **A-B.** Confocal images of the heart of *wild type* non injected control *tg(tp1:EGFP)*^*um14*^ embryo (A) and F0 *tg(tp1:EGFP)*^*um14*^ embryo injected with CRISPR/Cas9 against *kmt2d* (B). F0 injected embryos showed hypoplastic heart as seen in *kmt2d*^*zy59*^ null mutants (B). Notch positive cells (green) are mostly distributed in the atrio-ventricular valve of a 3dpf embryonic heart. Some endocardial cells in the ventricle and outflow tract are also observed (A). In F0 injected *kmt2d*-CRISPR mutants, hearts show a significant increase in the number of Notch positive cells in both ventricle and atrium. Ventral view of the heart, only middle sections of the whole dataset are shown. Scale bar = 25 µm. oft: out flow tract; ven, ventricle. **C.** Quantification of the amount of Notch positive cells in ventricle and atrium of control and injected embryos. N = 7 per group, unpaired two-tailed t-test, *p* = 0.0001 in ventricle t=4.95 dF=36, atrium *p* = 0.0001, t=10.99 dF= 36. **D.** RT-qPCR analysis of *wild type* sibling control embryos and *kmt2d*^*zy59*^ mutants for some components of the Notch signaling pathway. The Notch transcription factor *rbpja* was significantly up-regulated *kmt2d*^*zy59*^ embryos validating the results obtained in the F0 analysis. There was no significant difference found for *notch1b* and *hes1*. N = 4 per genotype with 2 technical replicates per gene and per genotype assessed; *elfα* was use as control gene. Ct values were normalized using *α-tubulin* as gene of reference; fold change of relative expression was calculated using the ΔΔCt method. Multiple t-test *p* < 0.0001 for *rbpja*, t=6.04, dF=24. **E-F.** Confocal images of *wild type* sibling control embryos (E) and *kmt2d*^*zy59*^ mutants (F) showing Rbpj protein expression levels and patterning at 5 dpf. Ventral view of the heart, only MIP of half dataset is shown. Scale bar = 25 µm. G. Summarized data and statistics from FACS experiment performed in 14 individual samples (7 *wild type* siblings and 7 *kmt2d* mutants). *Wild type tg(kdrl:GFP)* siblings (A) and *kmt2d*^*zy59*^;*tg(kdrl:GFP)* mutants were collected at 3 dpf, processed for IF, digested and prepared for FACS. Unpaired two-tailed t-test, % of EC cells Rbpj positives *p* = 0.63, n.s. t=0.49, dF=12; Rbpj MFI *p* = 0,0009, t=4.35 df=12.

Altogether our results show that Notch pathway is hyperactivated in endocardial cells of *kmt2d*^*zy59*^ mutants and demonstrate that this increased Notch activity is consequence of upregulated Notch pathway transcription factor Rbpj specifically in EC cells. To our knowledge, these results provide the first evidence of a regulatory link between Kmt2d and Notch signaling during physiological developmental processes in vertebrates.

### Pharmacological inhibition of Notch signaling rescues cardiovascular development in Kabuki Syndrome zebrafish mutants

Our results showed hyperactivation of Notch activity in endocardial and endothelial cells of *kmt2d* mutants, suggesting a molecular mechanism for the cardiovascular phenotype in KS. To test whether interference with this mechanism could rescue the cardiovascular phenotype in *kmt2d*^*zy59*^, inhibited the Notch pathway at the level of the Notch receptor cleavage by blocking y-secretase activity with DAPT (Fig. 7E) (58,61). *kmt2d*^*zy59*^;*tg(kdrl:GFP)* embryos and *wild type tg(kdrl:GFP)* siblings were treated with DAPT or DMSO (control group) from 1 dpf to 2 dpf, washed and assessed at 5dpf for cardiovascular development. *Wild-type* embryos treated with DAPT had other phenotypes predicted from inhibiting Notch as previously described (61): abnormal somite development, heart edema, disrupted vasculature (data not shown). Confocal images of *wild-type* DAPT treated embryos revealed aberrant heart morphology with hypoplastic ventricle and stretched cardiac tube likely as a consequence of pericardic edema (Fig. 7AB) evidenced by myosin staining of the sternohyoideus muscle (Fig. 7A-D asterisks, magenta labeling). Strikingly, DAPT treatment partially rescued aortic arch development and heart morphology in *kmt2d*^*zy59*^ mutant embryos (Fig. 7D). Heart ventricle volume was significantly increased in *kmt2d*^*zy59*^ DAPT treated embryos compared with the control groups: DMSO *kmt2d*^*zy59*^ embryos and *wild type* DAPT treated embryos (Fig. 7F; 2-way ANOVA multiple comparison test, n = 10 per condition, adjusted *p* values per group, see figure legend for *p value* details).

**Figure 7.**
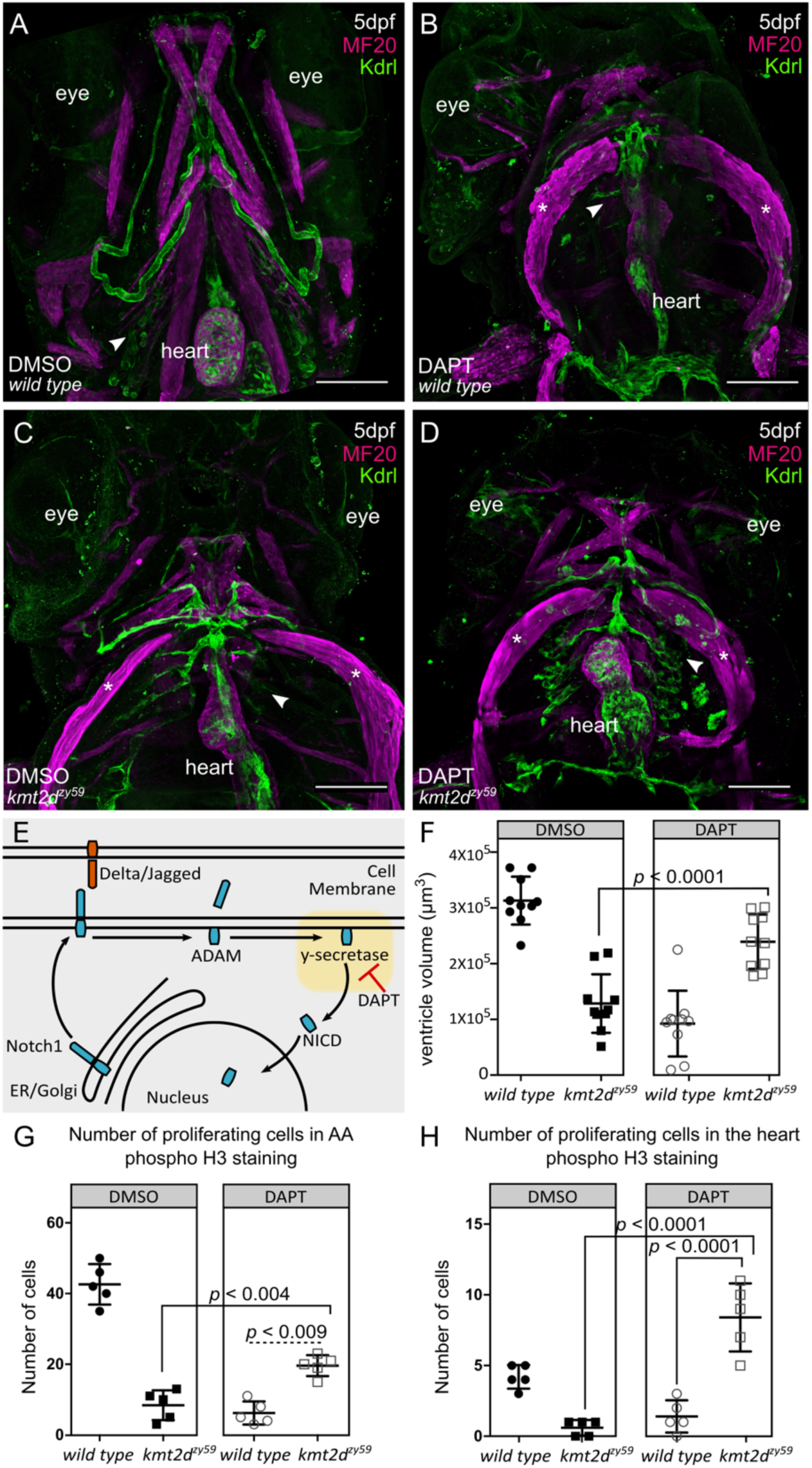
Inhibition of Notch pathway with DAPT rescues cardiovascular phenotype in *kmt2d* mutants by enhancing cell proliferation in endothelial and endocardial cells. **A-D.** Confocal images of *wild type* sibling (A, B) and *kmt2d*^*zy59*^ mutant (C, D) embryos at 5 dpf. DMSO as solvent control (A, C) and DAPT for Notch signaling inhibition (B, D) were applied to embryos of indicated genotypes. DAPT treatment affects cardiovascular development in *wild type* embryos (B) as a consequence of Notch signaling inhibition. *kmt2d*^*zy59*^ mutants that were treated with DAPT had rescued aortic arches development and partially rescued heart development (D) when compared with the *kmt2d*^*zy59*^ DMSO control group (C). Arrowheads indicate normal (A), disrupted (B,C) and rescued (D) aortic arches. Note that cardiovascular phenotype is rescued in DAPT-treated mutants despite of cardiac edema, as evidenced by the sternohyoideus deformation (asterisk; B, C, D, MF20 in magenta). **E.** Schematic of Notch signaling pathway showing DAPT inhibition of γ-secretase activity, the second cleavage step of Notch receptor processing. DAPT thus prevents NICD release to the cytoplasm for nuclear import. **F.** Cardiac ventricle volume measurements for DMSO control groups (*wild type* and *kmt2d*^*zy59*^) and DAPT treatment groups (*wild type* and *kmt2d*^*zy59*^). The volume of the ventricle chamber was significantly rescued in *kmt2d*^*zy59*^ mutant embryos after Notch pathway inhibition with DAPT. Measurements were blind and embryos were randomly selected from 3 different clutches. Genotype was confirmed after measurement by HRMA. 2-way ANOVA multiple comparison test adjusted *p* values per each condition as follow: *wild type* DMSO vs. wild type DAPT, *p* < 0.0001; *wild type* DMSO vs *kmt2d*^*zy59*^ DMSO, *p* < 0.0001; *wild type* DMSO vs. *kmt2d*^*zy59*^ DAPT, *p* = 0.0159; *wild type* DAPT vs *kmt2d*^*zy59*^ DMSO, *p = 0*.*5482*; *wild type* DAPT vs. *kmt2d*^*zy59*^ DAPT, *p* < 0.0001; *kmt2d*^*zy59*^ DMSO vs. *kmt2d*^*zy59*^ DAPT, *p* = 0.0001. **G.** Number of endothelial cells proliferating in the aortic arches region in each group treatment. There are significantly more proliferating endothelial cells in the aortic arches region of *kmt2d*^*zy59*^ mutant after DAPT treatment, indicating that the phenotype is rescued by increasing cell proliferation. 2-way ANOVA multiple comparison test adjusted *p* values per each condition as follow: *wild type* DMSO vs. wild type DAPT, *p* = 0.0001; *wild type* DMSO vs *kmt2d*^*zy59*^ DMSO, *p* = 0.0001; *wild type* DMSO vs. *kmt2d*^*zy59*^ DAPT, *p* = 0.0006; *wild type* DAPT vs *kmt2d*^*zy59*^ DMSO, *p = 0*.*6043*; *wild type* DAPT vs. *kmt2d*^*zy59*^DAPT, *p* = 0.0091; *kmt2d*^*zy59*^ DMSO vs. *kmt2d*^*zy59*^ DAPT, *p* = 0.0047. **H.** Number of endocardial cells proliferating in the heart per experimental group. There are significantly more proliferating endocardial cells in *kmt2d*^*zy59*^ mutant hearts after DAPT treatment, indicating that the phenotype is rescued by increasing endocardial cell proliferation. 2-way ANOVA multiple comparison test adjusted *p* values per each condition as follow: *wild type* DMSO vs. wild type DAPT, *p* = 0.0051; *wild type* DMSO vs *kmt2d*^*zy59*^ DMSO, *p* = 0.0307; *wild type* DMSO vs. *kmt2d*^*zy59*^ DAPT, *p* = 0.0013; *wild type* DAPT vs *kmt2d*^*zy59*^ DMSO, *p = 0*.*8106*; *wild type* DAPT vs. *kmt2d*^*zy59*^ DAPT, *p* < 0.0001; *kmt2d*^*zy59*^ DMSO vs. *kmt2d*^*zy59*^ DAPT, *p* < 0.0001. Cell count was blind and embryos were randomly selected from 2 different clutches (E, F). Genotype was confirmed after measurement by HRMA.

We then asked whether the rescue of aortic arches and heart development by Notch inhibition was due to a change in the proliferative capability of endothelial/endocardial cells. To test this, we analyzed cell proliferation by phospho S10 Histone 3 labeling and counted GFP positive cells (endothelial and endocardial cells) co-localizing with phospho S10 Histone 3 in a delineated area (Supp. Fig. 7A; white dashed line). Our analysis showed that DAPT treatment of *kmt2d* mutants induces a significant increase in the number of proliferating endothelial and endocardial cells in the aortic arches area and heart respectively.

## Discussion

Kabuki Syndrome is a rare multi-systemic developmental disorder mainly characterized by postnatal growth deficit, distinct facial features, hearing defects, abnormal neurologic development, immune dysfunction and congenital heart disease (CHD), predominantly left-sided defects and coarctation of the aorta (1,3,62). These cardinal features contribute to KS phenotypes with variable expressivity and different severity degrees. However, patient prognosis and morbidity mainly depends on early diagnosis and treatment of CHD and immune dysfunction (23). The mechanisms through which KMT2D mutations affects cardiovascular development remain unclear. Development and validation of a genetic KS animal model that provides a good platform for the analysis of cardiovascular phenotypes will allow a better understanding of the molecular mechanisms underlying the evolution of KMT2D-related CHD in a KS context.

In this study, we developed a genetic zebrafish model for Kabuki Syndrome that not only recapitulated cardinal phenotypic traits of the human pathology, but most importantly, allowed us to uncover previously unknown cardiovascular defects precipitated by abnormal endothelial/endocardial cell patterning. Through a combination of transcriptome analysis, F0 screening and FACS, we identified Notch signaling as a candidate pathway underlying the endothelial/endocardial phenotype. We also demonstrated that altered Notch pathway signaling was driven at the level of its nuclear transcription factor Rbpj. Importantly, drug inhibition of Notch signaling was able to rescue the cardiovascular phenotype in KS mutants (Fig. 8).

**Figure 8.**
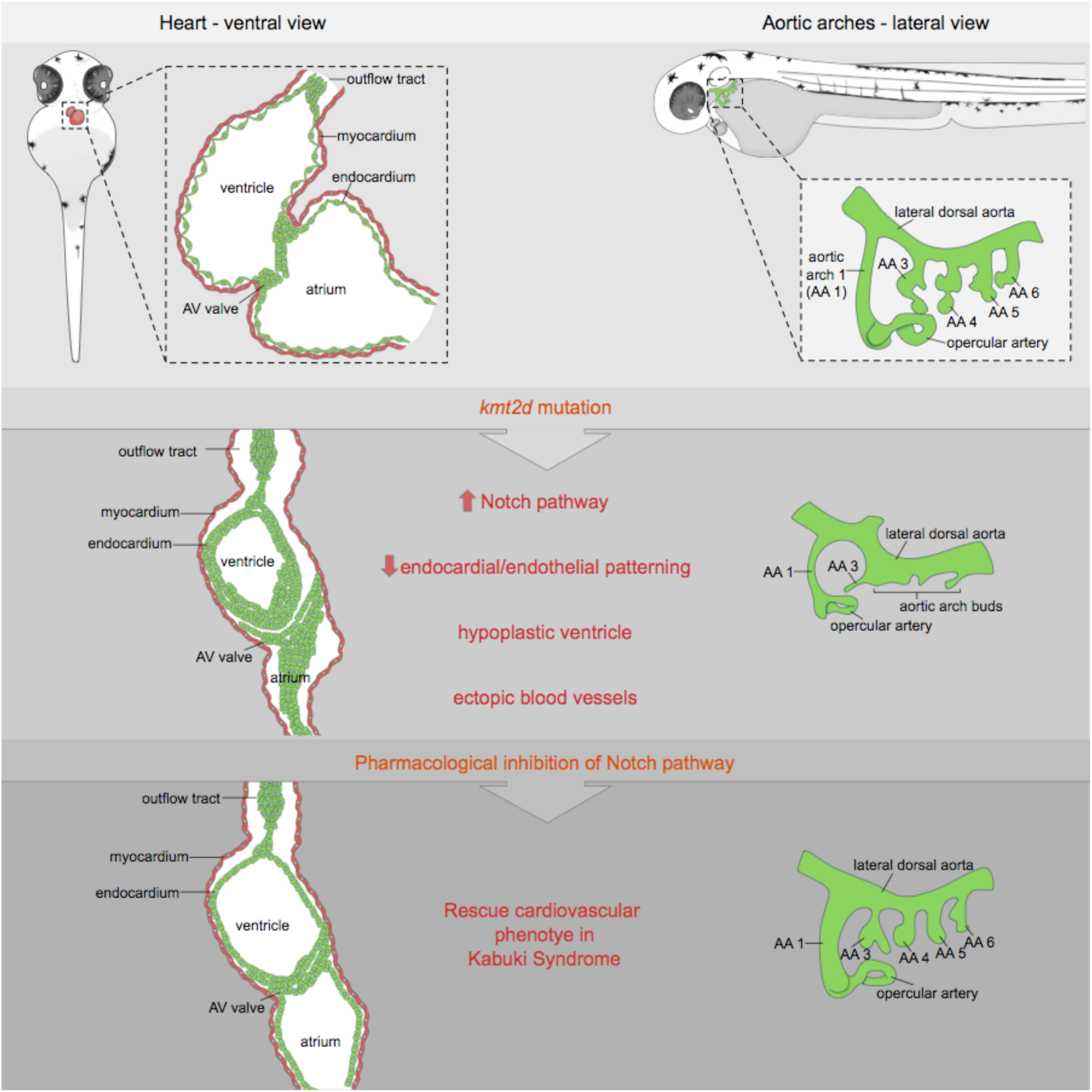
Model summary schematic. Schematic cartoon highlighting cardiovascular defects in our genetic zebrafish model for Kabuki Syndrome. Notch signaling is identified as primary candidate pathway underlying the endothelial/endocardial phenotype. Pharmacological inhibition of Notch signaling was able to rescue the cardiovascular phenotype in KS mutants.

Notch pathway is a known regulator of endocardium and vascular patterning during vertebrates development (54,63). Rbpj is the major transcriptional effector of Notch signaling (64). Recent studies in *Drosophila* demonstrated that KMT2D binds nuclear Notch co-activator complex (65), establishing a strong precedent of a regulatory link between Notch pathway and KMT2D (66). This report is consistent with our findings that loss of *kmt2d* produces a misbalance of Notch signaling at the level of transcription factor regulation. Our results demonstrate that Rbpj is upregulated at the transcript and protein level in zebrafish *kmt2d* mutants.

During canonical Notch signaling, ligand-receptor interactions of Notch components drives sequential proteolytic cleavage of the Notch receptor and nuclear translocation of the intracellular domain of Notch (NICD). Once there, NICD binds to the effector protein RBP-J and enables the induction of target genes by recruitment of co-activators in a cell-context dependent manner (67).

However, NICD cleavage also occurs independently of Notch ligand-receptor interaction; thereby providing basal levels of NICD that will bind Rbpj and regulate Notch signaling in a non-canonical fashion (68,69). In line with this paradigm, our results show that the canonical ligands and receptors of Notch pathway upstream of Rbpj are not affected in *kmt2d* mutants. Additionally, manipulating availability of NICD (canonical and non-canonical) by inhibiting its proteolytic cleavage with DAPT allowed us to rescue the *kmt2d* mutant cardiovascular phenotype. Altogether, these data indicate that the regulatory link between Kmt2d and Rbpj during cardiovascular development occurs via non-canonical Notch pathway in a ligand-independent manner. However, under the current paradigm, this hypothesis cannot entirely explain the full extent of our results. Even though the observed vascular mispatterning, ectopic angiogenesis and disorganized endocardium in our *kmt2d* mutants fits with a Notch pathway dysregulation, we do not assume that the underlying molecular mechanisms are the same in each scenario. Further studies are needed in order to fully understand the link of Kmt2d and non-canonical Notch pathway in the context of angiogenesis and vasculogenesis.

Pioneering studies demonstrated the importance of Kmt2d during myocardium development in mammals, but did not test a possible role in early endocardium morphogenesis (22). In this sense, our results in zebrafish *kmt2d* mutants demonstrate for first time a strong contribution of the endocardium to the hypoplastic ventricle phenotype described here. In contrast to our finding of a fundamental role of Kmt2d during endocardium patterning during zebrafish heart development, our myocardium analysis does not support a myocardial kmt2d mutant phenotype. Thus, we propose that the hypoplastic ventricle and overall cardiac phenotype in Kabuki Syndrome is mainly driven by endocardium mispatterning.

Current knowledge in Kmt2d function is focused on its roles as a H3K4 methyltrasferase (70). However, not much is known about Kmt2d non-histone substrates and gene regulatory networks. Our transcriptome data provides evidence for a broader functional spectrum of Kmt2d and offers strong gene candidates that we are pursuing in order to comprehend the multisystemic phenotype in zebrafish KS phenotype. Furthermore, our gene set enrichment analyses suggest that genes coding for core extracellular matrix (ECM) proteins have a strong early contribution in *kmt2d*^*zy59*^ mutant’s phenotype. This result suggests that the observed angiogenesis defect in *kmt2d*^*zy59*^ might be a consequence of an abnormal ECM, as previously demonstrated in other disease scenarios such as cancer development (71). Moreover, our results demonstrate that the observed endothelial/endocardial phenotype is not due to cell identity/specification failure nor hypoxia stress but a defect in the maintenance of cell behavior maintenance. Overall, these results indicate a novel regulatory link between Kmt2d and Notch pathway in cardiovascular patterning and suggest a possible therapeutic approach for ameliorating KS phenotypic traits of diseases caused by Kmt2d mutations.

## Materials and Methods

### Contact for reagent and resource sharing

Further information and requests for resources and reagents should be directed to and will be fulfilled by the Lead Contact, H. Joseph Yost

### Experimental model and subject details

Zebrafish embryos, larvae, and adults were produced, grown, and maintained according to standard protocols approved by the Institutional Animal Care and Use Committees of University of Utah. For experiments, zebrafish embryos ranging from 1 dpf to 7 dpf were used. Adults were maintained at ∼5 fish l^-1^ for all experiments in aquarium with controlled light cycle and water temperature at 28°C. Animals were fed 3 times a day.

Published strains used in this study include: wild-type AB, *tg(kdrl*:*EGFP)*^*la116*^ (72) and *Tg(EPV*.*Tp1-Mmu*.*Hbb:EGFP)*^*um14*^ referred in this manuscript as *tg(tp1:EGFP)*^*um14*^ (73).

## Method details

### Mutants Generation and genotyping

Mutants were generated in AB wild-type background. Mutagenesis was induced with CRISPR/Cas9 genome editing tools. Guide RNA was designed and synthesized at the University of Utah Mutation Generation and Detection Core Facility. Guide RNA was designed using as template *kmt2d* sequence available in Genome build Zv9, Ensembl annotation released version 79. Guide RNA target sequence corresponding to exon 8 is as following: 5’-GTATTGACTGTGGCATGCGA-3’. Genotyping was performed through High Resolution Melt Analysis (HRMA) (74) using CFX96 Touch Real-Time PCR Detection System. HRMA primer sequences are: forward, 5’—AGT TTT AGC GGT GCC GTG TG-3’; reverse, 5’-CCA CTG TTC AGA GCC AGG AAG-3’. Mutations were confirmed by DNA Sanger sequencing using the primers: forward, 5’-AGC TTG TAC AGA AGT TTG GCA A-3’; reverse, 5’-GGA AAA GTA CAC TTT AGA AAA CAG C-3’. Multiple alleles were isolated with ∼5 - 8 independent founders per allele. Mutation was confirmed and phenotypes analyzed in all of them. Lines are maintained as heterozygous in a *kdrl*:*EGFP* transgenic background.

### RNA extraction for RNA sequencing experiment

RNA was extracted from 50 individual embryos at 1 dpf obtained from a heterozygous × heterozygous(*kmt2d*^*+/zy59*^) cross from 2 parental crosses. We recorded parent id and order of processing for each embryo to be able to adjust for technical and biological variability. Genomic DNA and total RNA was extracted using Quick-DNA/RNA extraction kit from Zymo. After genotyping by HRMA, 6 embryos of each genotype (*wild type, heterozygous* and *mutant*) were selected, with 3 embryos from each of 2 parental crosses. A total of 18 samples were selected considering processing order and submitted to the High-Throughput Genomics and Bioinformatic Analysis Shared Resource at Huntsman Cancer Institute at the University of Utah. RNA concentrations per sample were on average 8 ng-μL and the RNA integrity Numbers (RIN) were 9 or higher. Libraries were built using Illumina TruSeq Stranded mRNA Library Preparation Kit with polyA selection and sequencing protocol was HiSeq 125 Cycle Paired-End Sequencing v4. The 18 samples were multiplexed in three sequencing lanes.

### Alcian Blue/Red alizarin and O-Dianisidine staining

Staining was performed at 4dpf as previously described (75). For O-Dianisidine experiment, 6 dpf embryos were fixed with 4% paraformaldehyde overnight. Fixed embryos were washed in PBS for three times and then incubated in the staining buffer (0.6 mg/mL o-dianisidine, 10 mM sodium acetate (pH 5.2), 0.65% hydrogen peroxide, and 40% ethanol) for 15 min in the dark.

### Immunofluorescence

Embryos were fixed over-night in 4% paraformaldehyde in PBS with triton 0.4%. Fixed embryos were washed with PBS 1x + triton 0.2% (PBST) 3 times for 15 minutes each in a rotator at room temperature. Pigment was removed incubating embryos for 15 minutes in a solution containing KOH 0.8%, Tween20 0.1%, H_2_O_2_ 0.9% and H_2_O to final volume; washed 3 times for 15 minutes each with PBST and incubated for 1 hour at room temperature with blocking solution (PBST + 5% goat serum). Primary antibody incubation was performed over night at 4°C. Primary antibody was removed and samples were washed with PBST 3 times for 30 minutes each. Samples were incubated with secondary antibody over night at 4°C. Antibody dilutions were performed in PBST with 1% goat serum. After secondary antibody incubation, samples were washed with PBST 3 times for 30 minutes each and rinsed for 15 minutes with increasing concentrations of glycerol (30%, 50% and 80%) in PBS 1X. Stained samples were mounted in low melt agarose at 1% for confocal imaging.

Primary antibody concentration was established following manufacturer guidelines; secondary antibody concentration was 1:500 in all cases. Antibodies are detailed in the Key Resource Table. Anti-GFP antibody was used to enhance endogenous transgenic fluorescence detection in *tg(kdrl:GFP)* line. Nuclear staining is not shown in processed images but it was performed for cell quantification experiments. DAPI was used in a concentration of 1mg.ml^-1^ and applied during secondary antibody incubation step.

### Single embryo fluorescent activated cell sorting (FACS)

Embryos were fixed and processed for immunofluorescence as described in the previous section. After secondary antibody incubation and washes, samples were distributed placing 1 single embryo per 1.5mL tube. PBST was completely removed and 1mL of pre-warmed TrypLE (1X) solution was added in each tube. Samples were incubated for 1.5 hour at 37°C. Pipetting every 10 minutes was performed to aid in dissociation. At the end of 1.5 hours, solution was checked for cell aggregates. TrypLE reaction was stopped by adding CaCl_2_ to a final concentration of 1mM and fetal bovine serum to 10%. Each sample was filtered through a 40μm nylon mesh filter (Fisher 22363547) into a 50mL tube. Filtered samples were spun down at 2,500 RPM for 10 minutes at 4°C. Supernatant was discarded and cells of each individual embryo were resuspended in 300μL of PBS 1X.

For Cell cycle profiling, cells were resuspended in DAPI staining solution (Triton 0.1%, PBS 1X, DAPI 1mg.ml^-1^). For accurate sample gating, controls in each experiment included: samples without fluorescence (for excluding yolk autofluorescence), samples without fluorescence with secondary antibody incubation only (for excluding secondary antibody unspecificity), samples with DAPI, samples with fluorescence of interest and samples with DAPI and fluorescence of interest. Samples were processed for flow cytometry analysis on a BD FACS Canto and Propel Labs Avalon Cell Sorter at the University of Utah Flow Cytometry Facility. FACS data was analyzed with FlowJo (v9) software.

### Confocal images acquisition, processing and analysis

Images were acquired in Zeiss LSM880 with Airyscan fast module following optimal parameters for super resolution (SR) mode acquisition. Image acquisition was performed keeping the same laser parameters for the channel of interests (channel to be measured/quantified). Acquisition parameters corresponding to context markers were enhanced as required for image clarity. Output files were processed for Airyscan processing with Zen Black software keeping default parameters. Output files (.czi) were processed with Fiji (ImageJ) to build maximum intensity projections for slides of interests or the complete data set.

Imaris software (v9.2) was used to reconstruct 3D microscopy images, measure heart ventricle volume, cell count and cell morphology numbers. The Imaris *surfaces* feature was used to analyze the volume of the ventricle by generating a surface in the lumen of each ventricle. Endocardial cells were used as limit so the space delimited by them was considered ventricle lumen and measured. Cell numbers were gathered by creating a surface containing the channel corresponding to myocardium, endocardium or Notch positive cells and the channel corresponding to DAPI was masked in each tissue/cell specific surface. The *measurement points* tool was used to count number of nuclei in each surface.

Myocardial cells area and circularity were measured using the *surfaces* tool in manual mode. Alcama antibody was used a reference marker for myocardial cell limits. The *clipping plane* feature of Imaris was used to visualize each cell and validate the surface created. Details of this method can be found in Supplementary figure 3 legend.

### Time-lapse experiments

Time-lapse experiments were performed in embryos from a *kmt2d*^*zy59*^;*tg(kdrl:GFP)* het by het cross at 2 dpf, 3dpf and 4 dpf. In each case, genotype was confirmed at the end of the experiment by phenotype screening and HRMA. Samples were placed in embryo water with N-Phenylthiourea 1X (PTU) for avoiding pigment formation and anaesthetized with MS-222 (150mg.L^-1^). Samples were embedded in 0.6% low melt agarose with MS-222 in embryo water.

Acquisition was performed in Zeiss LSM880 with Airyscan fast module using 10x dry objective and default *optimum* parameters. Parameters of acquired data as follow: 16bit depth image, 1.8 zoom, 1176×1176px image size and minimum z-stack interval for capturing development of aortic arches (approx. 467.56 × 467.56 × 2.88μm). Images were acquired every 20 minutes for a length of 12 hours. After acquisition period, samples were recovered for genotyping and/or immunofluorescence and SR confocal imaging.

### F0 experiment

For analyzing the effects of *kmt2d* mutation in a Notch reporter background, the *kmt2* guide RNA/Cas9 (described in the section *Mutant Generation*) was injected in *tg(tp1:EGFP)*^*um14*^ embryos. Approximately 100 embryos per clutch were injected and 40 of them were processed for *kmt2d* mutation validation though HRMA. Injected embryos showed a range of phenotype from mild to strong effects. A total of 7 embryos per condition (injected – not injected) were selected based on their gross morphology similarity with class II *kmt2d*^*zy59*^ mutants (Figure 1B).

### Quantitative PCR experiment

3dpf wild type/heterozygous and *kmt2*^*zy59*^ mutant embryos were lysed in TRI Reagent (Zymo) and processed for RNA using a Zymo Quick-RNA kit. The total RNA was used for cDNA synthesis with BioRad iScript 5X master mix. Subsequent cDNA was used for qPCR reactions (CFX96 Touc Real-Time PCR Detection System) with the following gene primers: *notch1b, rbpja, hes1 and elfα*. Primer and probe sequences are listed in the Key Resources table. Delta Ct calculations were normalized to *elfα* expression levels and control sibling expression levels. The number of samples used was N=4 per genotype with 2 technical replicates per gene and per genotype assessed.

### Pharmacological treatment

For Notch pathway inhibition, embryos from a *kmt2d*^*zy59*^;*tg(kdrl:GFP)* het by het cross were treated with 50μM DAPT from 1dpf to 2dpf. After treatment, embryos were washed with fresh embryo water and maintained until 5dpf for immunofluorescence sample processing. After fixation, DAPT treated embryos were tail-clipped for DNA extraction and genotypification by HRMA. All treatments, including controls were as following: *wild type* DMSO, *wild type* DAPT, *kmt2d*^*zy59*^ DMSO and *kmt2d*^*zy59*^ DAPT. For hypoxia induction, *tg(kdrl:GFP)* and *kmt2d*^*zy59*^*;tg(kdrl:GFP)* embryos were treated from 3 dpf to 4 dpf with 100μM DMOG (HIF-Hydroxylase Inhibitor). After treatment, embryos were washed and processed for IF and confocal imaging. DMSO was used as control for drug solvent in mutant and wild-type genotypes.

#### Quantification and statistical analysis

### General Statistical analysis

Sample sizes were chosen based on previous publications and are defined in each figure legend. No sample was excluded from the analysis. Unless explicitly expressed in the text or figure legend (Figure 7FH, Supplementary Figure 3C), the experiments were not randomized and the investigators were not blinded to allocation during experiments and outcome assessment. All statistical values are displayed as mean ± standard deviation with exception of Supplementary Figure 3AC were 25^th^ percentile, 50th percentile and 75th percentile are shown. Sample sizes, statistical test and *p* values are indicated in the figures or figure legends. Statistical significance was assigned at *p* < 0.05. Statistical tests were performed using Prism 7 software and R package.

### RNA differential expression analysis

Transcriptome was aligned with STAR (76) to Genome build Zv9, Ensembl annotation released version 79. In this study, differential expression analysis was performed between 6 wild-type versus 6 mutant samples with DESeq2 (77) using the negative binomial likelihood ratio test. Heterozygous samples were excluded from this analysis. An initial cutoff of 5% false discovery rate was set.

### Gene set enrichment analysis (GSEA)

To perform GSEA analysis we used the differential expression analysis gene list ranked by *p* values. We then filtered the list to obtain genes that have one-to-one orthology to human genes as specified in the Ensembl Compara database that were retrieved with BiomaRt package (78). Gene sets obtained from the Molecular Signatures Database (MSigDB). The resulting list was additionally filtered using a mean normalized count cutoff of 5. The number of resulting genes identifiers analyzed was 9128 out of 33737. GSEA pre-ranked analysis (10,000,000 permutations, minimum term size of 15, maximum term size of 500) was then performed using fgsea package (https://github.com/ctlab/fgsea/)(79)

The analysis was performed independently with 8 selected gene sets and subsets from MSigDB: H: hallmark gene sets, CP: Canonical pathways, CP: KEGG: KEGG gene sets, CP:REACTOME: Reactome gene sets, BP: GO biological process, CC: GO cellular component, MF: GO molecular function, C7: immunologic signatures. Results cutoff was set at 5% FDR and plotted using R programming interface.

## Supporting information

## Data Availability

RNA sequencing data used for building Figure 1D is available in Supplementary Table 1. Complete results from gene set enrichment analysis are available in Supplementary table 2. Raw data is available at https://b2b.hci.utah.edu/gnomex/gnomexFlex.jsp?requestNumber=475R

## Acknowledgements

We thank Dr. Saulius Sumanas, Dr. Nathan Lawson and Dr. Neil Chi for providing reagents or transgenic fish lines.

## Author Contributions

Conceptualization, M.A.S., M.T.F. and H.J.Y.

Methodology, M.A.S. M.T.F. and H.J.Y.

Software, B.L.D.

Formal Analysis, M.A.S., B.L.D.and T.T.P.H.

Investigation, M.A.S. and T.T.P.H.

Resources, M.T.F. and H.J.Y.

Writing – Original Draft, M.A.S.

Writing – Review & Editing, M.A.S., B.L.D., M.T.F and H.J.Y.

Visualization, M.A.S. and B.L.D.

Diagrams and cartoons – M.A.S.

Funding Acquisition, M.A.S. and H.J.Y.

## Supplemental Information Titles and Legends

**Supplementary Figure 1:**
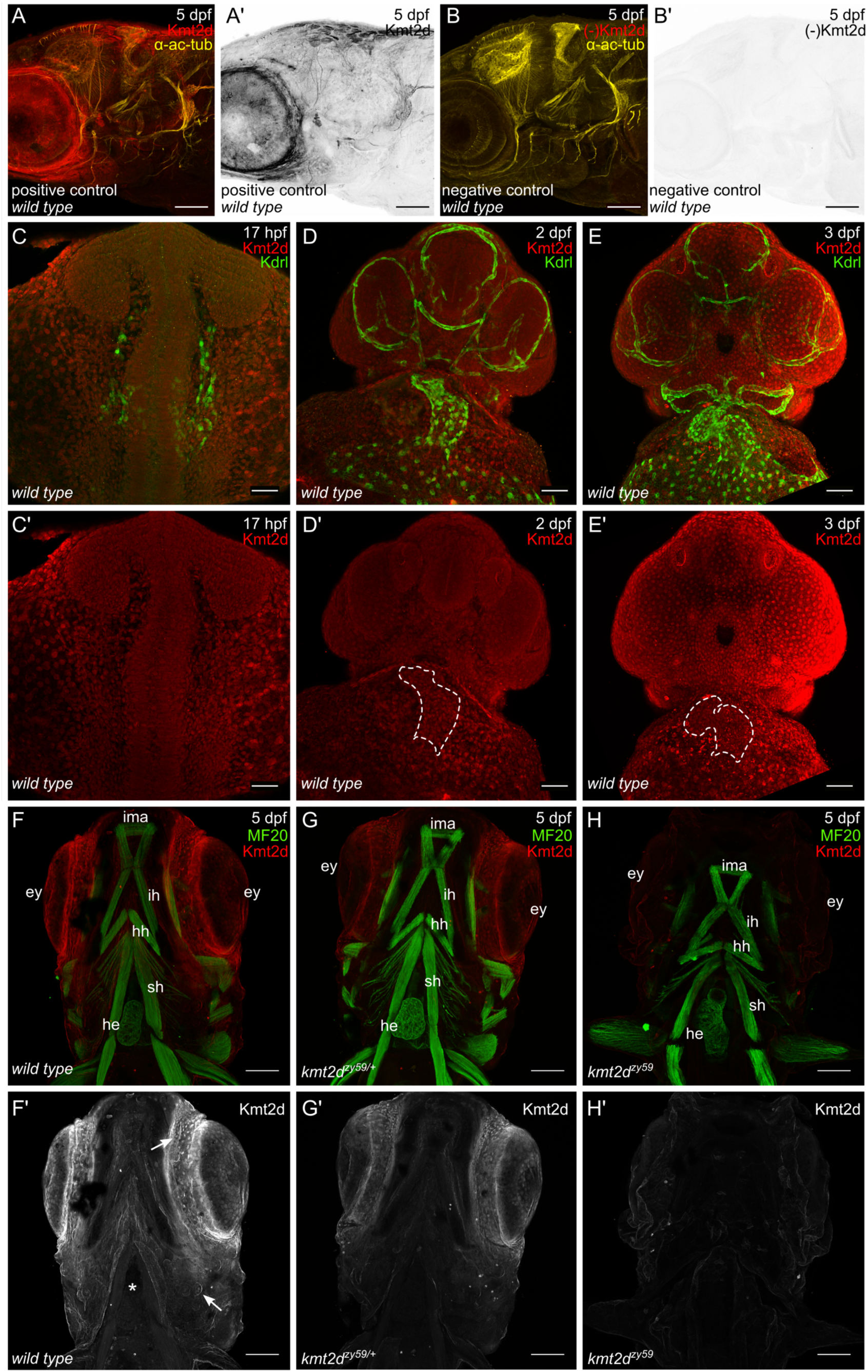
Kmt2d protein expression. A-B. Kmt2d antibody technical control. Confocal images of 5 dpf zebrafish embryo in a lateral view. Immunofluorescence was performed against Kmt2d (red and black) and alpha acetylated tubulin (*α*-ac-tub, yellow) as context marker. Kmt2d antibody specificity for zebrafish was assessed through Immunofluorescence negative control. (A) Positive control of Kmt2d Antibody. (B) Kmt2d negative control. Samples were processed in parallel to the positive controls. Primary antibody incubation was omitted and samples were incubated with secondary antibodies. (A’-B’) Kmt2d channel was selected, set as gray scale and look up table was inverted in order to enhance contrast. C-E. Kmt2d protein expression time course. Confocal images of ventral views of zebrafish *tg(kdrl:GFP)* embryos at 17 hpf (C-C’’), 2 dpf (D-D’’) and 3 dpf (E-E’’). Immunofluorescence was performed against Kmt2d (red) and GFP (Kdrl, green) as context marker. (C, D and E) Merge for Kmt2d and Kdrl. (C’, D’ and E’) Channel for Kmt2d (red). White dashed line delineates the heart (D’ and E’). Images were processed as MIP. F-H. Kmt2d null mutant validation. Confocal images of 5 dpf zebrafish embryos in a ventral view. Images were processed as maximum intensity projections (MIP) Immunofluorescence was performed against Kmt2d (red and black) and myosin heavy chain (MF20, green) as context marker. Samples were genotyped by HRMA after image acquisition. (F) Homozygous *wild type*, (G) heterozygous and (H) homozygous mutant. Note the lack of Kmt2 protein expression confirming *kmt2d*^*zy59*^ as null mutant. (F’-H’) Kmt2d channel was selected, set as gray scale and look up table was inverted in order to enhance contrast.

**Supplementary Figure 2:**
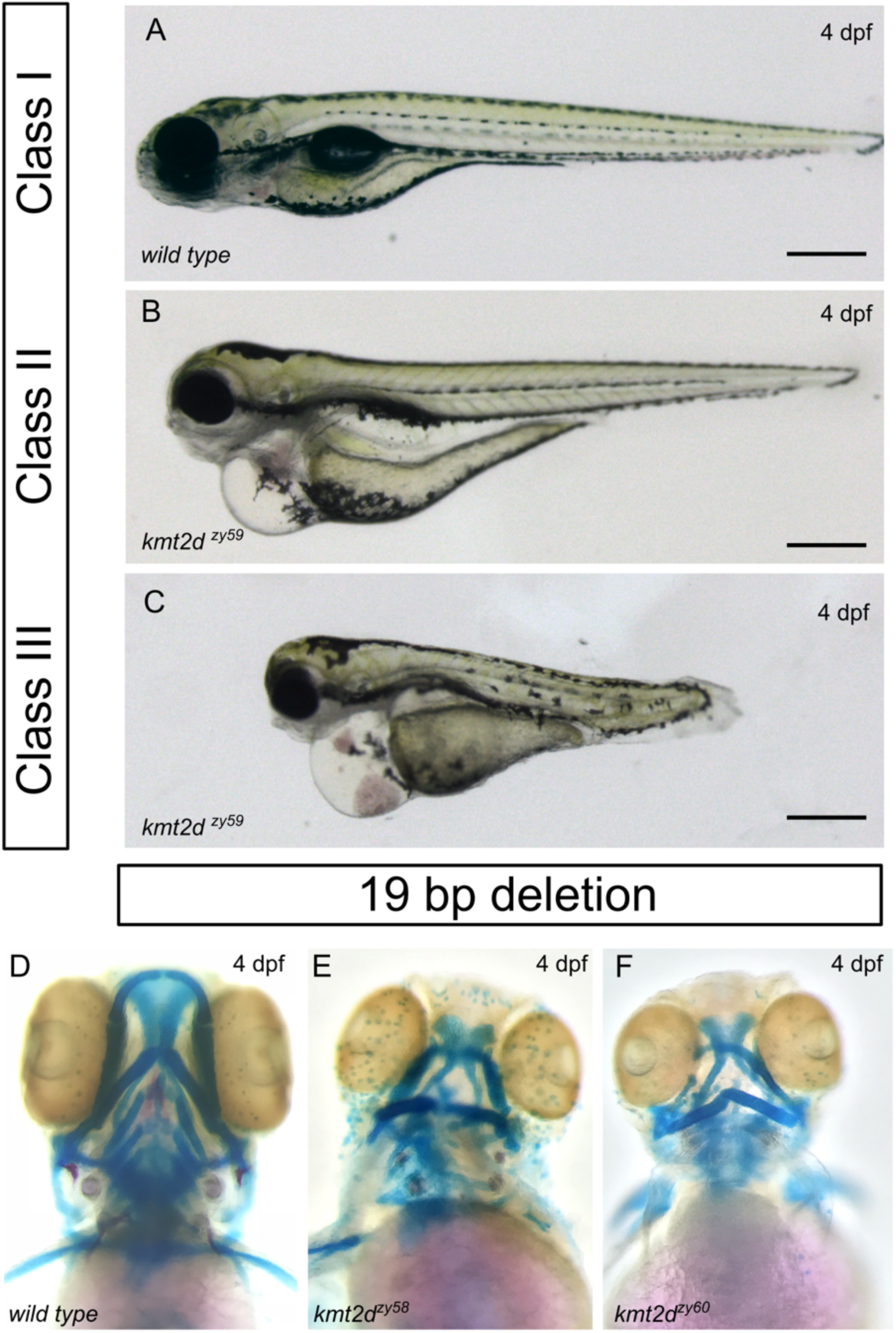
*kmt2d* mutant phenotype at 4dpf. A-C. Lateral view of zebrafish *wild type* sibling embryo (A) and *kmt2d*^*zy59*^ mutants (B, C) at 4dpf. At 4dpf *kmt2d*^*zy59*^ embryos develop general body edema that increases gradually at later stages (not shown). D-F. Alcian blue/ Alizarin red staining in two additional mutant alleles

**Supplementary Figure 3:**
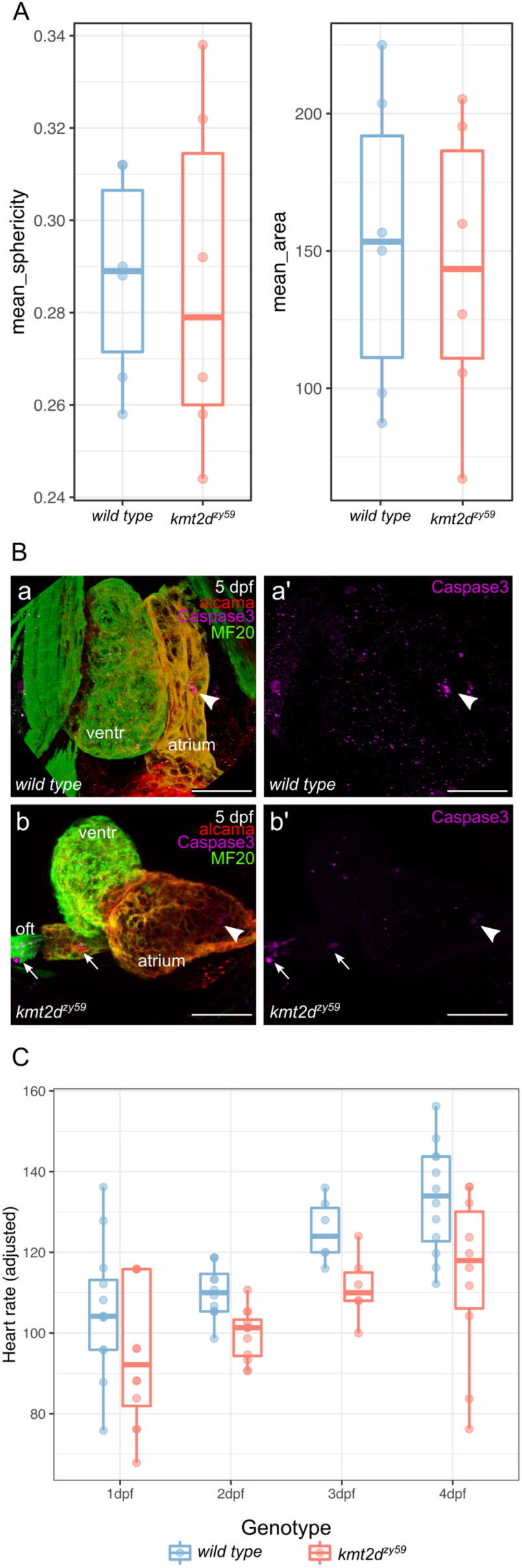
Analysis of myocardial cell morphology, apoptosis and heart rate, in *wild type* siblings and *kmt2d*^*zy59*^ mutants. A. Myocardial cell shape analysis in *kmt2d*^*zy59*^ mutants at 3 dpf. *Wild type* sibling and *kmt2d*^*zy59*^ mutant embryos were processed for immunofluoresce against Alcama for cell-cell boundaries and myosine heavy chain (MF20) for myocardium context. Z-stacks were analyzed with Imaris software. Area and circularity were measured in 5 different cells from the outer curvature of the ventricle. Averaged values are plotted. There is no significant difference in cardiomyocytes shape in wild type samples vs. mutants. t-test, *p* < 0.583 n.s. t=0.59 dF=5 for area and *p* < 0.946 n.s. t=0.71 dF=5 for circularity. B. Apoptosis analysis in *wild type* vs. *kmt2d*^*zy59*^ mutant heart. Confocal images of *wild type* sibling and *kmt2d*^*zy59*^ at 5 dpf. The heart was acquired from a ventral view. Immunofluorescence was performed against active-caspase3 for apoptosis evaluation and Alcama and MF20 as context markers. Arrows and arrowheads point to apoptotic cells. C. Heart Rate comparison in *wild type* siblings vs. *kmt2d*^*zy59*^ mutants at 1, 2, 3 and 4 dpf. Embryos were placed individually in a 96 well plate. Measurements were performed at each time point to the same animal subject every time in a blind fashion until day 3-4 where the phenotype was apparent. Heart beat count was performed for 15 seconds without anesthetic to avoid any secondary effects that could impact heart rate. Heart rate values were adjusted according to the ANOVA model, for both experiment and time points variability *p* value = 0.000264, F (1,76) = 14.647.

**Supplementary Figure 4:**
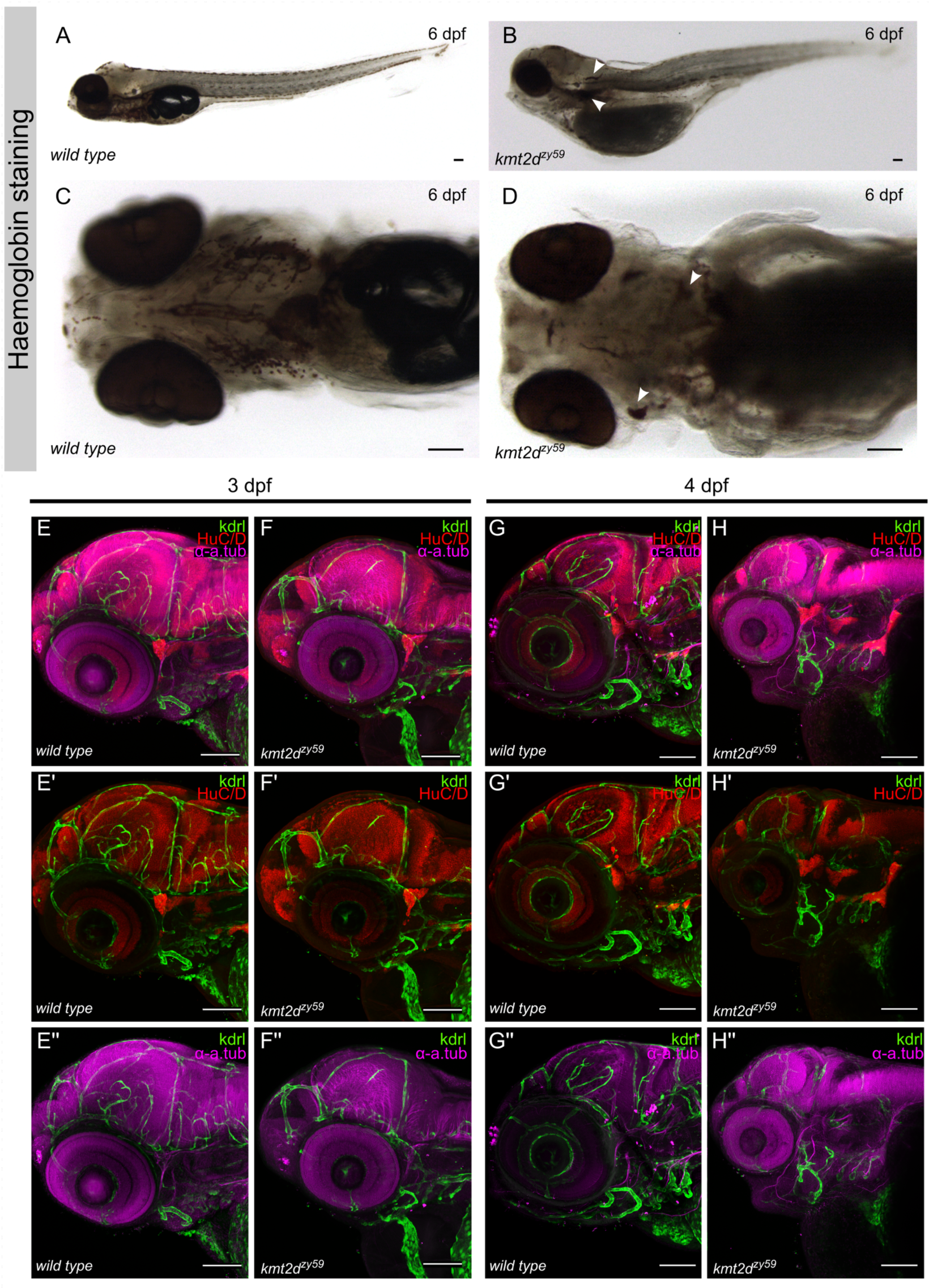
Vascular network analysis in *wild type* siblings and *kmt2d*^*zy59*^ mutants. A-D. *o-dianisidine* staining for assessing vasculature integrity in *kmt2d*^*zy59*^ and *wild type* siblings at 6 dpf. Lateral views (A, B) and cranial-ventral views (C, D) of *wild type* sibling (A, C) and *kmt2d*^*zy59*^ mutant (B, D) at 6 dpf. White arrowheads indicate blood aggregates in the region of AA and head. Scale bar = 100 µm E-H. Vascular development at 3 dpf and 4 dpf in *wild type* sibling vs. *kmt2d*^*zy59*^ mutant embryos. Confocal images of cranio-lateral views at 3 dpf (E, F) and 4 dpf (G, H) in *wild type* (E-E’’, G, G’’) and mutant (F-F’’, H, H’’) embryos. Immunofluorescence was performed against GFP, for enhancing Kdrl signal, HuC/D and *α*-acetylated tubulin as context markers.

**Supplementary Figure 5:**
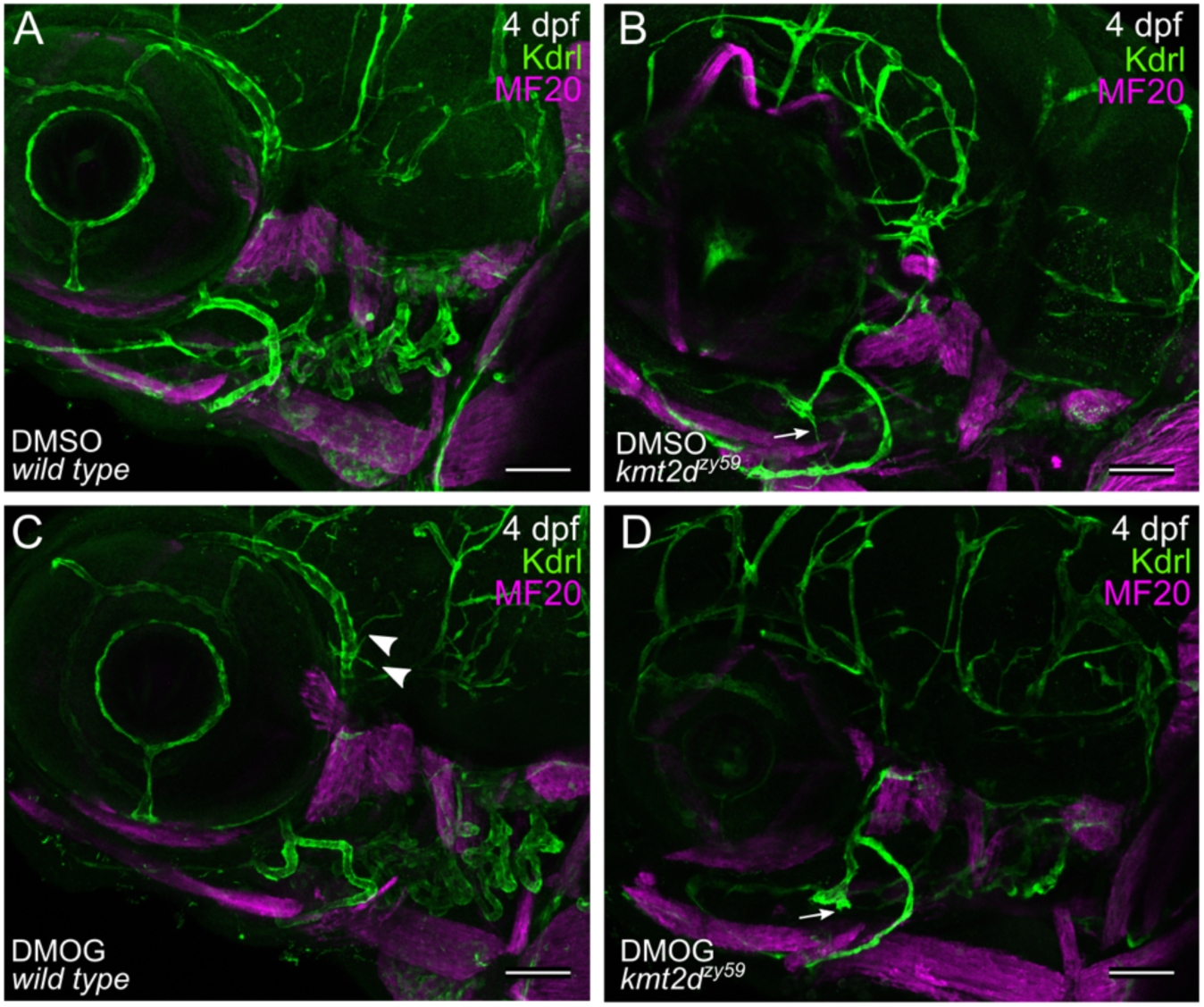
Vascular development in induced hypoxia conditions in *wild type* and *kmt2d*^*zy59*^ embryos. Confocal images show cranial-lateral view of vasculature in *wild type tg(kdrl:GFP)* sibling (A) and *kmt2d*^*zy59*^;*tg(kdrl:GFP)* mutants at 4 dpf. A-B.DMSO controls for both, wild type sibling and *kmt2d* mutant. C-D. DMOG treated embryos. Treatment was performed from 3 to 4 dpf. White arrowheads indicate hypoxia-induced blood vessel sprouting. White arrows (B and D) indicate *kmt2d* mutation-dependent ectopic blood vessel formation in both DMSO control and DMOG treated embryos.

**Supplementary Figure 6:**
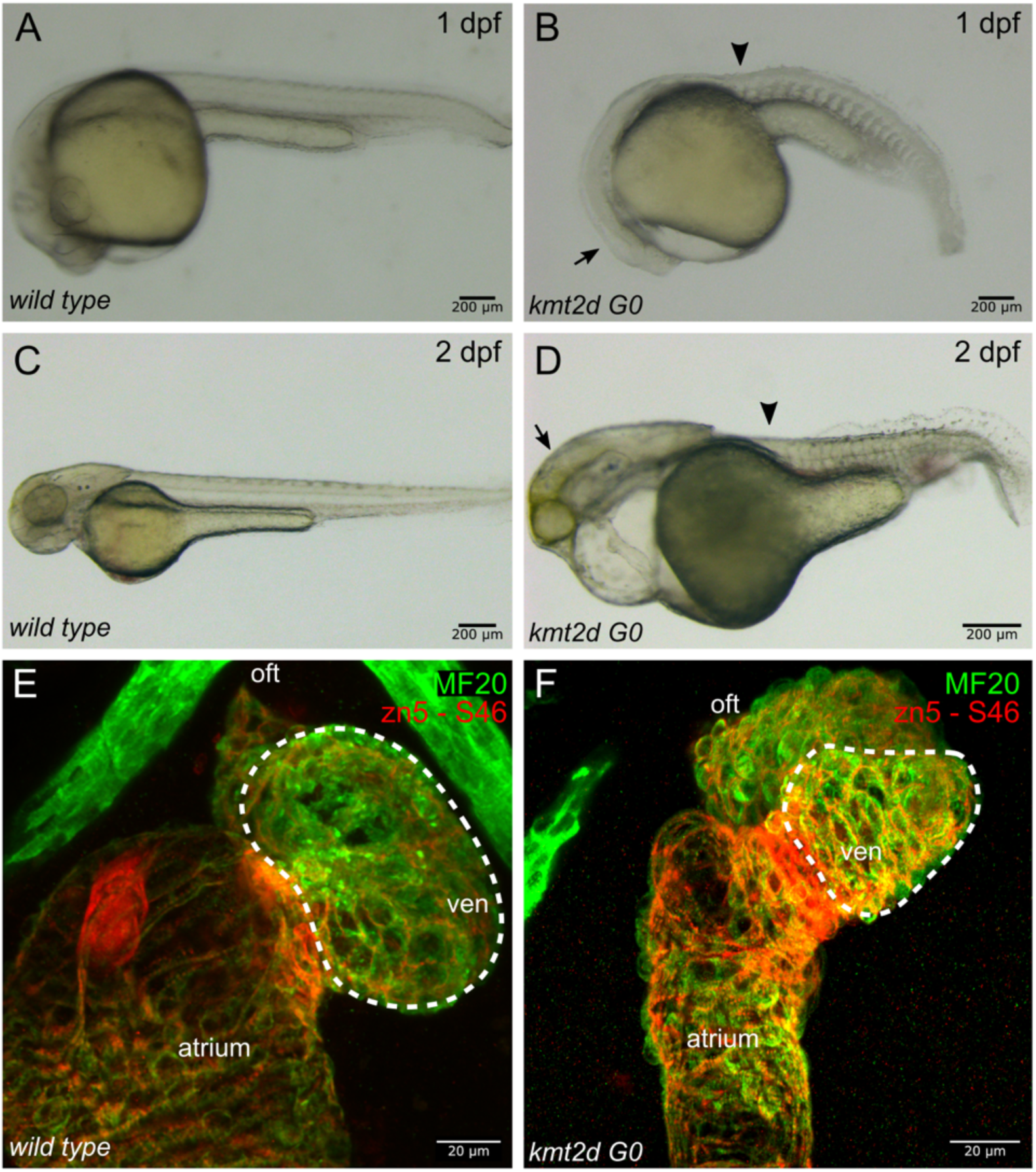
F0 phenotype validation. CRISPR/Cas9 injection against kmt2d produces comparable phenotype to the observed in germline mutants (arrows and arrowheads). A, C, E, Non injected controls. B, D, F, injected embryos. E, F, confocal images of non injected controls and kmt2d injected embryos. IF was performed for Myosin heavy chain (M20, green), Alcama (zn5, red) and Myosin heavy chain, atrium specific (S46, red) as general myocardium morphology markers. Dashed white line highlights hypoplastic heart as a consequence of mutated *kmt2d* through CRISPR injection.

**Supplementary Figure 7:**
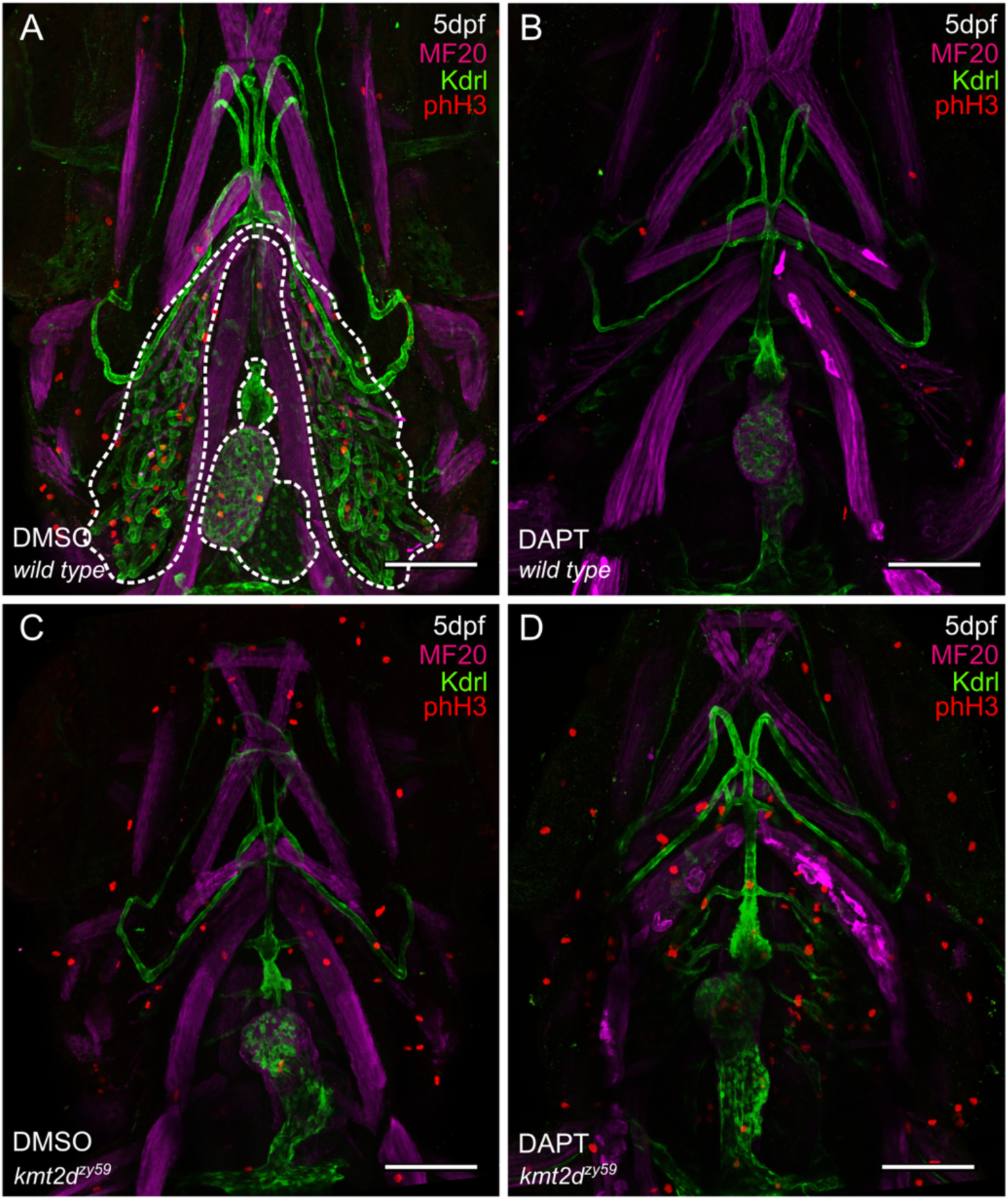
Proliferation assay for validating drug rescue phenotype. **A-D**. Confocal images of *wild type* sibling (A, B) and *kmt2d*^*zy59*^ mutant (C, D) embryos at 5 dpf. DMSO as solvent control (A, C) and DAPT for Notch signaling inhibition (B, D) were applied to embryos of indicated genotypes. Immunofluorescense against GFP was perform to enhance Kdrl:GFP signal (endothelium and endocardium). MF20 (myosin) was use as context marker for muscle. phH3 (cell proliferation) marks mitotic cells. Note the increased phH3 signal in cardiovascular area in *kmt2d* mutants after DAPT treatment (D).

### Supplementary Materials (Videos)

Video 1: 3D render of cardiovascular patterning of *wild type* sibling at 5dpf.

Video 2: 3D render of cardiovascular patterning of *kmt2d*^*zy59*^ mutant at 5dpf

Videos 3 and 4: 20 embryos from a *kmt2d*^+/-^ by *kmt2d*^+/-^ cross were mounted in 0.6% agarose for overnight time lapse imaging. Lateral view with head towards viewer’s left. Genotypes corresponding to *wild type/heterozygous* or *mutants* were assigned at the end of the experiment at 3dpf when phenotype was evident. One wild type embryo (Video 3) and one *kmt2d*^*zy59*^ mutant embryo (Video 4) are shown as examples. Other supplementary videos are available on request. Note AA 3-6 develops and extends almost overlapping with ORA in *wild type* embryo (Video 3). The AA growth occurs in a cranio-caudal direction with AA3 development before AA6. In contrast, *kmt2d* mutant embryo does not develop proper AA primary sprouting. Endothelial cell sprouting and extension is hyperactive, particularly in the more ventral-caudal region of the LDA, were AA6 should develop. Moreover, AA6 shows an abnormal primary sprout before the rest of AA (Video 4).

## Resources Table

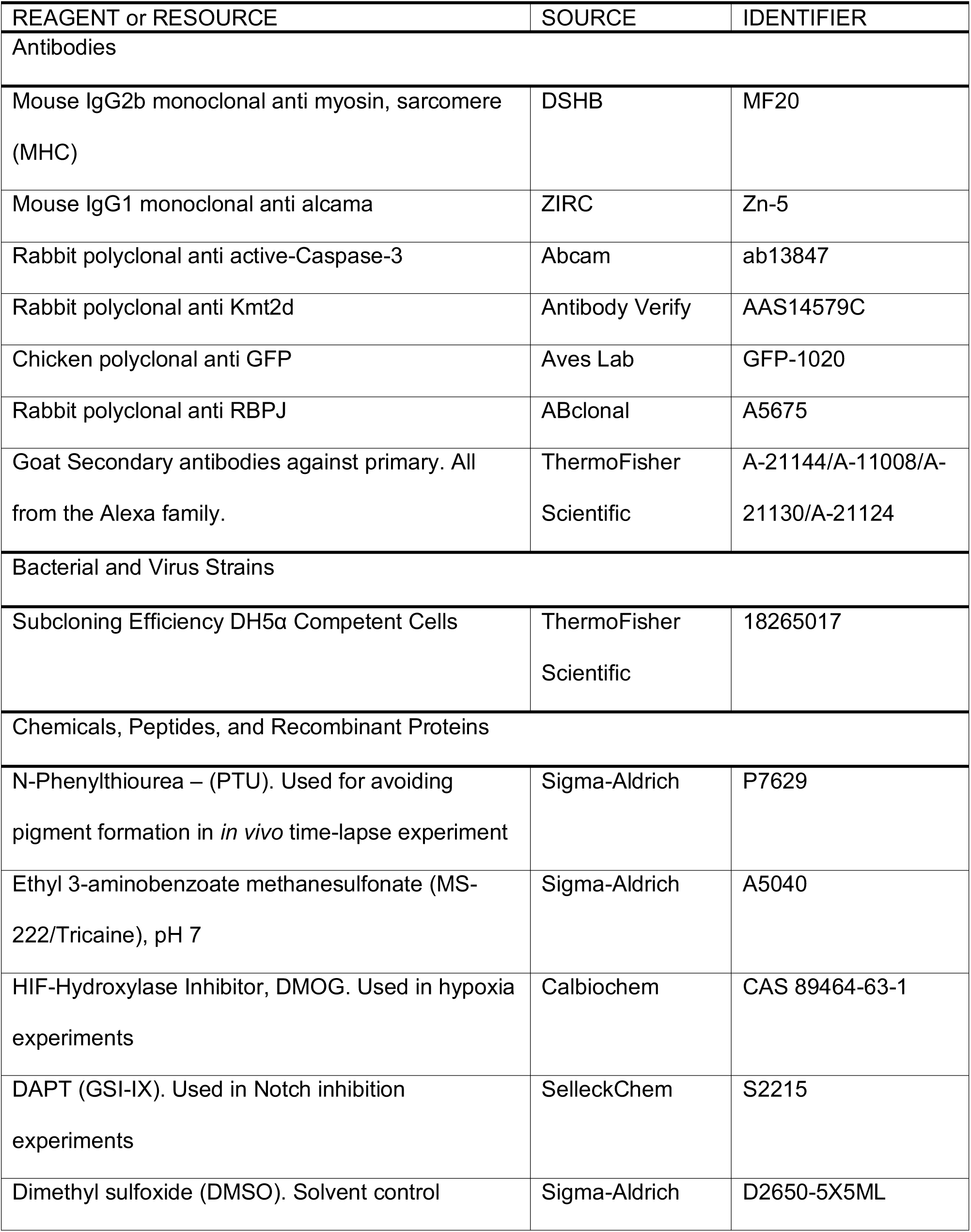
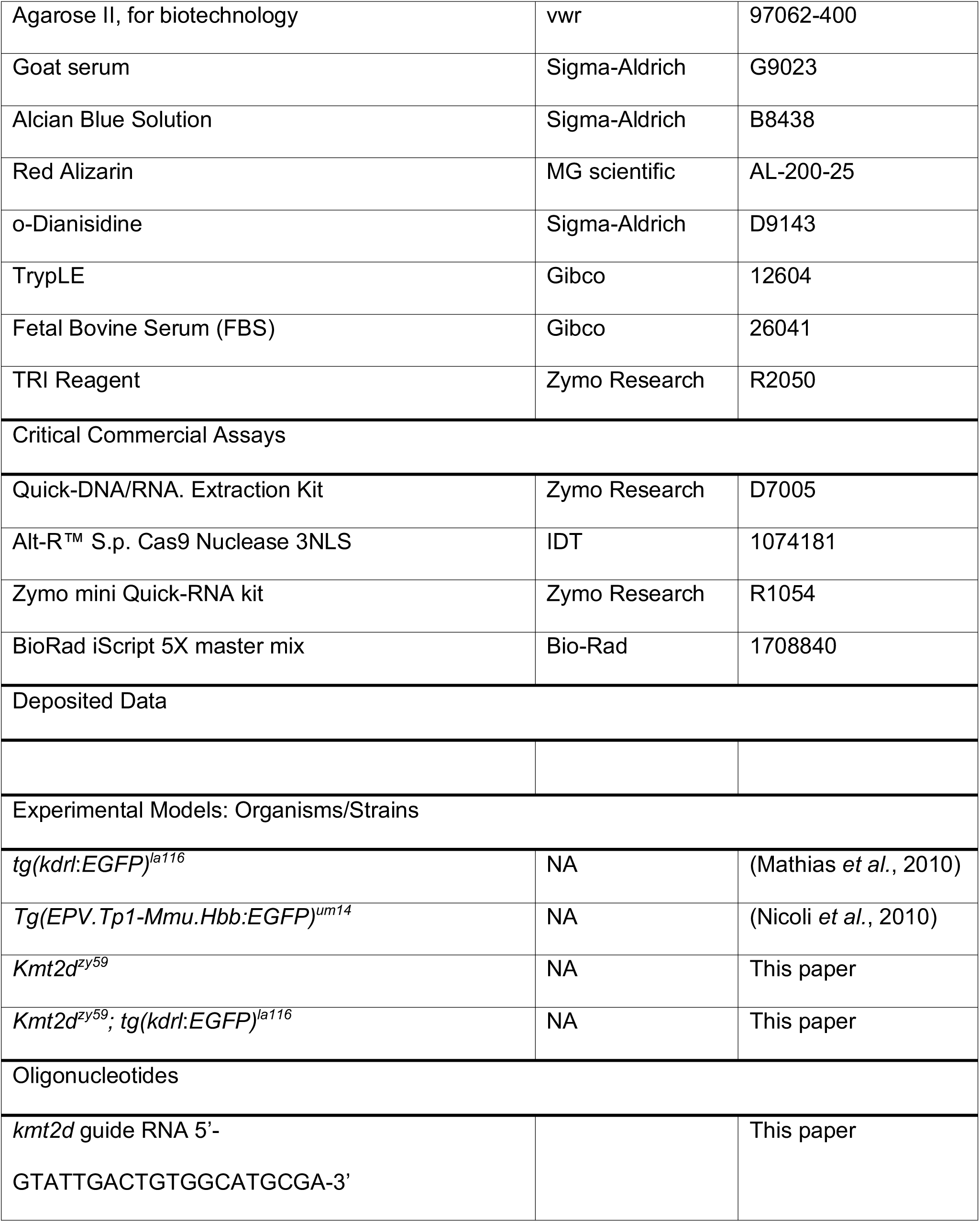
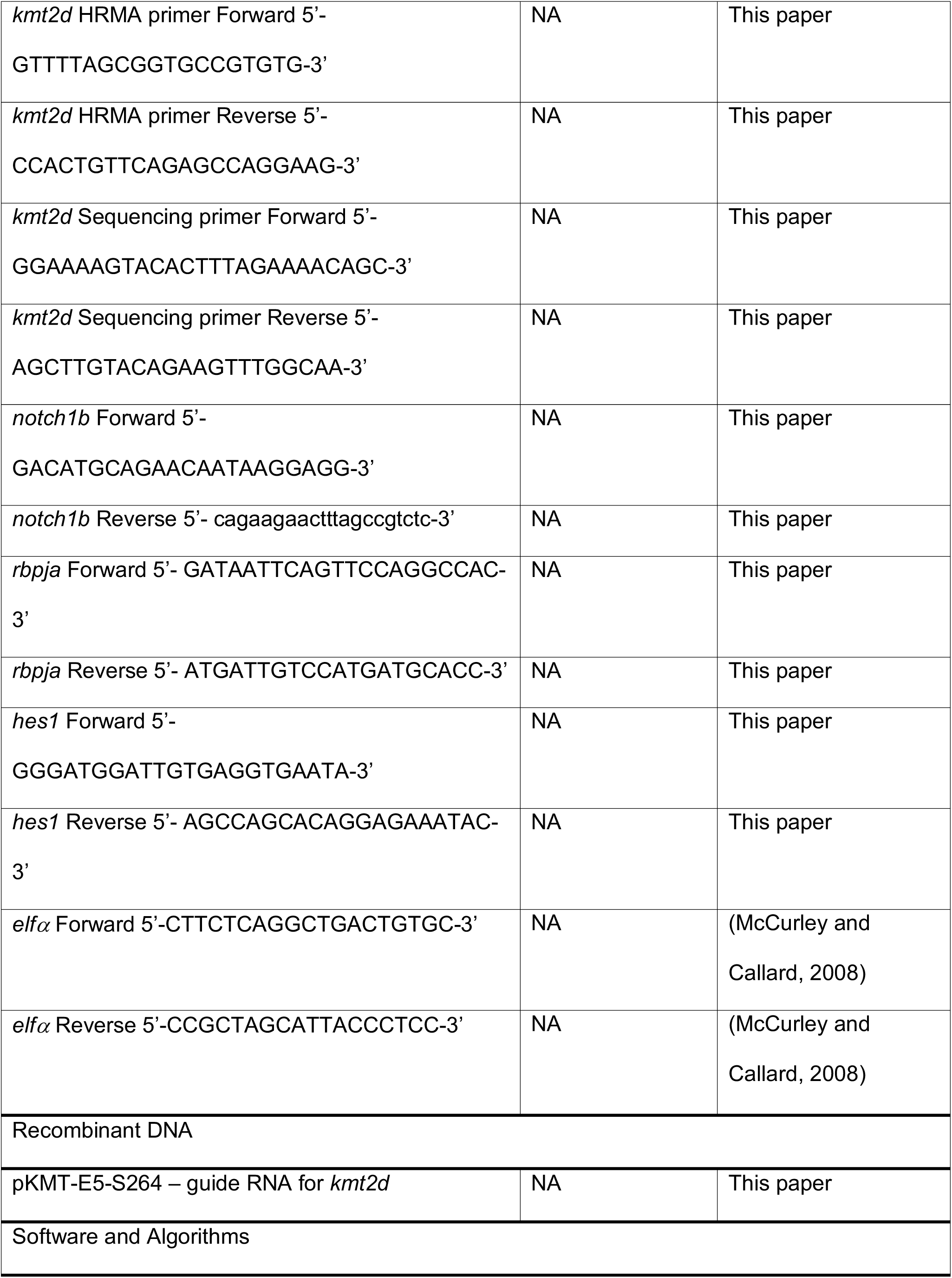
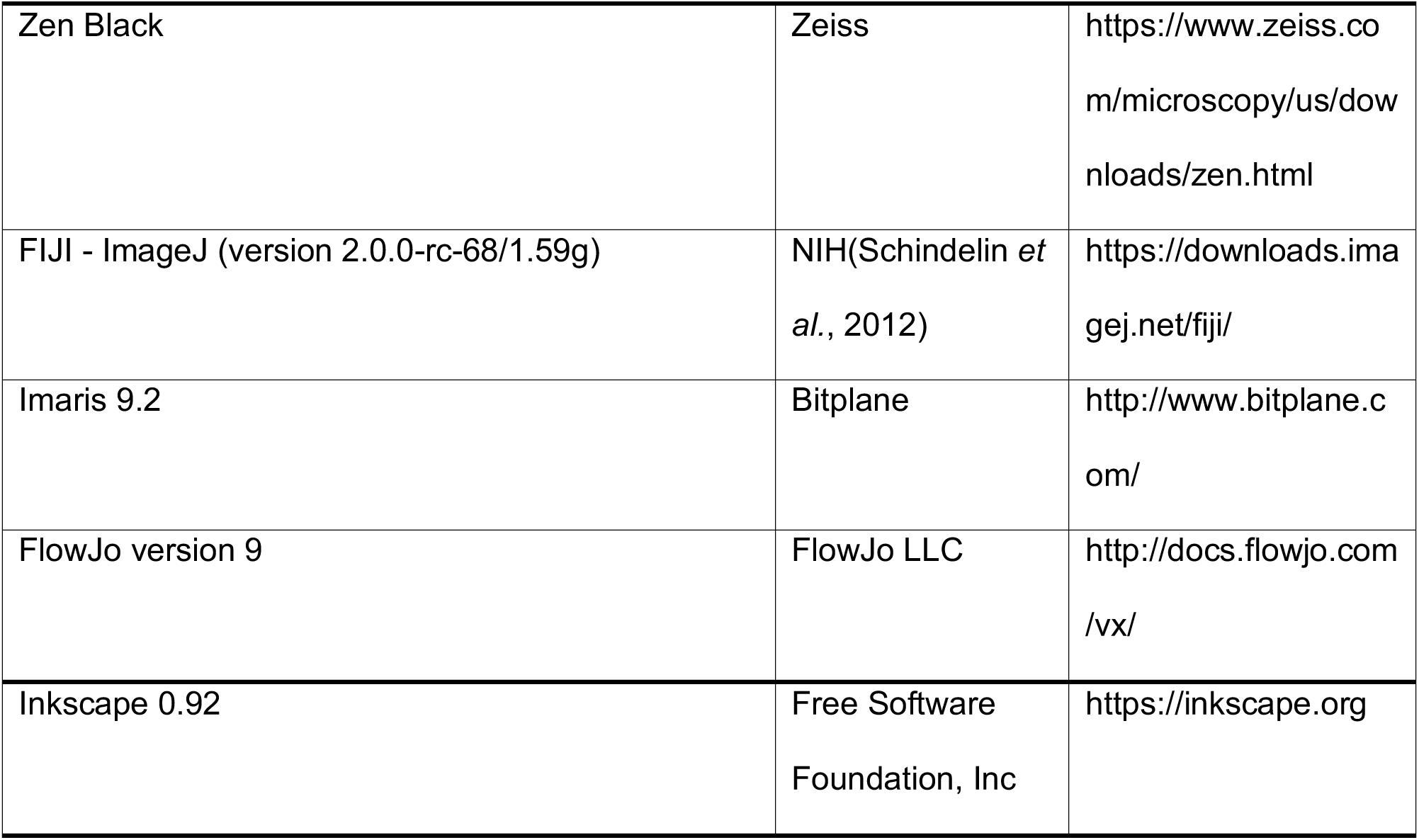

