## Supplementary material for "Inhibition of Notch signaling rescues cardiovascular development in Kabuki Syndrome"

### Supplementary Tables with Titles and Legends

#### Supplementary Table 1: Text-mining of top 50 differentially regulated genes

Top 50 gene candidate at a 5% FDR (base mean, log2 fold change and *p*-adjusted values are specified). Manual text-mining was performed for each candidate. Categories were established considering available information for cell compartment, biological function, region of expression and mammal orthologues data.

| ensembl_gene_id | external_gene_name | baseMean | log2FoldChange | padj | region_expression | category | source | link | |
|---|---|---|---|---|---|---|---|---|---|
| **ENSDARG00000018264** | trim101 | 223.266953 | -3.4205734 | 4.92E-25 | skeletal_muscle | muscle | zfin | http://zfin.org/ZDB-GENE-040801-100 | |
| **ENSDARG00000039929** | ckmt2b | 68.3279066 | -1.2735683 | 1.79E-05 | skeletal_muscle | muscle | zfin | http://zfin.org/ZDB-GENE-040426-1654 | |
| **ENSDARG00000061827** | zbtb4 | 141.480592 | -1.0032933 | 4.16E-09 | mesoderm | muscle | zfin | https://zfin.org/ZDB-GENE-100318-5 | |
| **ENSDARG00000075842** | pigt | 61.6656557 | -0.8932759 | 1.33E-05 | somite | muscle | zfin | https://zfin.org/ZDB-GENE-090313-46 | |
| **ENSDARG00000039265** | arhgap4a | 38.0616206 | -1.8567648 | 1.18E-08 | neural_tissue | neural | zfin - OMIN | http://zfin.org/ZDB-GENE-040426-1229 | https://omim.org/entry/300023?search=arhgap4&highlight=arhgap4 |
| **ENSDARG00000094426** | her4.2 | 692.157419 | -1.4117474 | 1.23E-12 | neural_tissue | neural | zfin | http://zfin.org/ZDB-GENE-060815-1 | |
| **ENSDARG00000010347** | acer1 | 112.087892 | -1.2215564 | 9.54E-07 | neural_tissue | neural | zfin | http://zfin.org/ZDB-GENE-050417-70 | |
| **ENSDARG00000070770** | her4.3 | 564.221021 | -1.1805975 | 4.19E-05 | neural_tissue | neural | zfin | https://zfin.org/ZDB-GENE-081030-7 | |
| **ENSDARG00000037140** | pfkfb1 | 214.42707 | -0.8844084 | 1.19E-11 | neural_tissue | neural | zfin | https://zfin.org/ZDB-GENE-030131-5664 | |
| **ENSDARG00000034896** | ldb2b | 250.609608 | -0.6198509 | 1.33E-05 | neural_tissue | neural | zfin | https://zfin.org/ZDB-GENE-990415-137 | |
| **ENSDARG00000033533** | CCDC115 | 524.303483 | -0.4989453 | 3.29E-05 | neural_tissue | neural | zfin - OMIN | https://zfin.org/ZDB-GENE-050227-20 | https://omim.org/entry/613734 |
| **ENSDARG00000069981** | cspg5a | 829.926292 | -0.4933447 | 1.04E-05 | neural_tissue | neural | zfin - OMIN | https://zfin.org/ZDB-GENE-080425-4 | https://omim.org/entry/606775 |
| **ENSDARG00000023174** | fez1 | 524.997667 | -0.4801264 | 1.54E-06 | neural_tissue | neural | zfin | https://zfin.org/ZDB-GENE-040426-2723 | |

| | | | | | | | | | |
|---|---|---|---|---|---|---|---|---|---|
| **ENSDARG00000076081** | sncgb | 353.493441 | -0.4567478 | 2.63E-05 | neural_tissue | neural | zfin | https://zfin.org/ZDB-GENE-050522-235 | |
| **ENSDARG00000030547** | rnd1 | 448.576717 | 0.90929143 | 6.37E-08 | neural_tissue | neural | zfin | https://zfin.org/ZDB-GENE-040630-6 | |
| **ENSDARG00000089549** | BAALC | 88.3687299 | 1.28939661 | 1.03E-09 | neural_tissue | neural | zfin - OMIN | https://zfin.org/ZDB-GENE-081022-24 | https://omim.org/entry/606602 |
| **ENSDARG00000090472** | ttll10 | 138.533229 | 1.96887323 | 7.30E-23 | neural_tissue | neural | zfin | https://zfin.org/ZDB-GENE-081104-350 | |
| **ENSDARG00000091235** | CABZ01015525.1 | 47.5085214 | -3.4152642 | 4.01E-05 | vascular_tissue | neural and cardiovascular | Institute of Cardiovascular Regeneration, Goethe University Frankfurt | http://angiogenes.uni-frankfurt.de/transcript?search=&rows_per_page_int=25&page_number=546 | |
| **ENSDARG00000054817** | ppp1r14c | 5.54292909 | -5.4382032 | 2.69E-05 | neural_cardiac_tissue | neural and cardiovascular | OMIN | https://omim.org/entry/613242 | |
| **ENSDARG00000020788** | sla2 | 23.4693679 | -2.3799544 | 9.45E-11 | immune_system | neural and cardiovascular | zfin - OMIN | http://zfin.org/ZDB-GENE-080204-98 | https://omim.org/entry/606577 |
| **ENSDARG00000089920** | CU571255.1 | 47.1329575 | -1.2207048 | 5.22E-06 | cardiac_skeletal_muscle | neural and cardiovascular | zfin | https://zfin.org/ZDB-GENE-041111-277 | |
| **ENSDARG00000070486** | rbp7b | 985.092104 | -0.9525541 | 1.57E-06 | neural_cardiac_muscle_tissue | neural and cardiovascular | zfin | https://zfin.org/ZDB-GENE-081022-134 | |
| **ENSDARG00000002644** | rgs5a | 164.568151 | -0.6718394 | 2.06E-06 | vascular_tissue | neural and cardiovascular | zfin | https://zfin.org/ZDB-GENE-030131-7570 | |
| **ENSDARG00000056831** | gng2 | 366.318584 | -0.5695759 | 1.79E-05 | neural_vascular_tissue | neural and cardiovascular | zfin | https://zfin.org/ZDB-GENE-050417-59 | |
| **ENSDARG00000086826** | sult6b1 | 1349.51741 | -0.5633785 | 1.35E-12 | neural_vascular_tissue | neural and cardiovascular | zfin | https://zfin.org/ZDB-GENE-050417-228 | |
| **ENSDARG00000056499** | ca6 | 1780.87737 | -0.5397003 | 2.46E-09 | neural_vascular_tissue | neural and cardiovascular | zfin | https://zfin.org/ZDB-GENE-030131-7091 | |

| | | | | | | | | | |
|---|---|---|---|---|---|---|---|---|---|
| **ENSDARG00000013855** | slc12a3 | 493.854667 | -0.5100772 | 5.31E-06 | neural_cardiac_gut | neural and cardiovascular | zfin | https://zfin.org/ZDB-GENE-030131-9505 | |
| **ENSDARG00000020785** | lama4 | 1142.44679 | -0.4982175 | 3.09E-05 | cardiac_vascular_tissue | neural and cardiovascular | zfin | https://zfin.org/ZDB-GENE-040724-213 | |
| **ENSDARG00000063538** | kalrnb | 505.638601 | -0.4433807 | 1.31E-05 | blood | neural and cardiovascular | zfin | https://zfin.org/ZDB-GENE-060421-7244 | |
| **ENSDARG00000056929** | kdm6bb | 1769.32634 | -0.4381307 | 1.16E-06 | neural_cardiac_vascular_tissue | neural and cardiovascular | zfin | https://zfin.org/ZDB-GENE-040724-166 | |
| **ENSDARG00000056075** | rca2.1 | 1170.46545 | -0.3794535 | 1.24E-06 | neural_cardiac_gut | neural and cardiovascular | zfin | https://zfin.org/ZDB-GENE-050320-60 | |
| **ENSDARG00000018404** | krt18 | 16821.7388 | 0.53661864 | 3.32E-05 | cardiac_vascular__tissue_pharyngeal_arch | neural and cardiovascular | zfin | https://zfin.org/ZDB-GENE-030411-6 | |
| **ENSDARG00000095512** | rca2.2 | 43.3431095 | 1.47416929 | 3.04E-09 | neural_cardiac_liver_tissue | neural and cardiovascular | zfin | https://zfin.org/ZDB-GENE-060503-646 | |
| **ENSDARG00000002945** | bgnb | 49.9919093 | 1.76077257 | 1.11E-09 | neural_cardiac_vascular_tissue | neural and cardiovascular | zfin - OMIN | https://zfin.org/ZDB-GENE-040426-21 | https://omim.org/entry/300989 |
| **ENSDARG00000004836** | dnajc5ab | 1423.26869 | 0.39387467 | 8.17E-09 | neural_testis_tissue | neural and reproductive | zfin - OMIN | https://zfin.org/ZDB-GENE-081021-2 | https://omim.org/entry/611203?search=dnajc5&highlight=dnajc5 |
| **ENSDARG00000056515** | spsb1 | 570.752435 | 0.53790646 | 2.71E-08 | neural_muscle_ovary_testis | neural and reproductive | zfin | https://zfin.org/ZDB-GENE-030131-6122 | |
| **ENSDARG00000005464** | dnase1l3 | 710.501878 | -0.7709107 | 1.53E-05 | neural_tissue_blood_pharyngeal_arch | neural, blood and pharyngeal arches | zfin | https://zfin.org/ZDB-GENE-040808-35 | |
| **ENSDARG00000089489** | CR762484.3 | 272.882325 | -0.6017719 | 3.93E-10 | neural_tissue_hematopoietic | neural, blood and | zfin | https://zfin.org/ZDB-GENE-000804-1 | |

| | | | | | | | | | |
|---|---|---|---|---|---|---|---|---|---|
| | | | | | | pharyng eal arches | | | |
| **ENSDARG000 00057863** | dnmt5 | 924.31 5763 | 0.3627808 9 | 1.3 1E-05 | neural_tissue_blood_phary ngeal_arch | neural, blood and pharyng eal arches | zfin | https://zfin.org/ZDB-GENE-050314-2 | |
| **ENSDARG000 00057426** | oard1 | 162.83 978 | - 2.2705516 | 5.4 1E-39 | | other | zfin | http://zfin.org/ZDB-GENE-050522-480 | |
| **ENSDARG000 00052336** | ociad2 | 98.655 3524 | - 1.0976163 | 3.9 5E-09 | | other | zfin | https://zfin.org/ZDB-GENE-041014-253 | |
| **ENSDARG000 00034714** | esyt1a | 381.76 1581 | - 0.7463954 | 8.3 7E-08 | spleen_adipose_tissue | other | zfin - OMIN | https://zfin.org/ZDB-GENE-090311-55 | https://omim.org/entry/616670 |
| **ENSDARG000 00070657** | pa2g4b | 7279.5 0801 | - 0.5600499 | 1.6 4E-06 | | other | zfin | https://zfin.org/ZDB-GENE-030131-2182 | |
| **ENSDARG000 00037071** | rps26 | 8956.1 6534 | 0.3952103 7 | 4.6 5E-06 | | other | zfin | https://zfin.org/ZDB-GENE-030131-8606 | |
| **ENSDARG000 00056122** | gdi1 | 2743.8 4085 | 0.5841122 7 | 4.4 1E-12 | | other | zfin | https://zfin.org/ZDB-GENE-050522-504 | |
| **ENSDARG000 00017773** | slc16a12a | 153.26 096 | 0.9022992 5 | 1.6 4E-06 | | other | zfin | https://zfin.org/ZDB-GENE-080721-24 | |

**Supplementary Table 2: Gene set enrichment analysis.**
Analysis was performed by converting zebrafish gene names to human gene names using exclusively genes with a one-to-one ortholog relationship. The number of resulting genes identifiers analyzed was 9128 out of 33737 (Genome build Zv9, Ensembl annotation released version 79). Adjusted p-values were calculated per category. Normalized Enrichment Score (NES) of gene sets with a FDR of 5% (blue dots) and 15% (purple dots) were plotted to summarized GSEA results.

| pathway | pval | padj | ES | NES | nMoreExtreme | size | leadingEdge | gene_set |
|---|---|---|---|---|---|---|---|---|
| REACTOME_INWARDLY_RECTIFYING_K_CHANNELS | 0.00183887 | 0.14651897 | 0.60694807 | 1.7259953 | 17947 | 16 | ABCC9\|KCNJ14\|KCNJ9\|GNG3\|GNG8\|KCNJ15\|GNB2\|KCNJ8\|GNG12\|GNB1\|GNG5 | CP_REACTOME |
| HALLMARK_PANCREAS_BETA_CELLS | 0.00259443 | 0.12453248 | 0.54881544 | 1.64856683 | 25705 | 22 | PCSK2\|ISL1\|NKX6-1\|HNF1A\|FOXA2\|SRPRB\|SPCS1\|SYT13\|SEC11A\|CHGA\|SST\|SRP14\|SLC2A2 | HALLMARK |
| REACTOME_G_ALPHA_Z_SIGNALLING_EVENTS | 0.00602768 | 0.14651897 | 0.56309388 | 1.61932356 | 59043 | 17 | RGS19\|ADRA2A\|RGS4\|GNG3\|PRKCQ\|GNG8\|GNB2\|GNG12\|GNB1\|GNAS\|GNG5 | CP_REACTOME |
| REACTOME_REGULATION_OF_WATER_BALANCE_BY_RENAL_AQUAPORINS | 0.00815677 | 0.147657 | 0.54544515 | 1.58469989 | 80143 | 18 | GNG3\|AQP4\|PRKAR2B\|GNG8\|PRKACG\|GNB2\|GNG12\|GNB1\|RAB11A\|GNAS\|GNG5\|ADCY8\|PRKAR1B\|GNG7\|RAB11FIP2\|ADCY5 | CP_REACTOME |
| REACTOME_VIF_MEDIATED_DEGRADATION_OF_APOBEC3G | 0.00182778 | 0.14651897 | 0.49247548 | 1.58275467 | 18258 | 36 | PSMB10\|PSMB6\|PSMC2\|PSMC4\|PSMB3\|PSMB5\|RBX1\|PSMD8\|PSMB4\|PSMA3\|PSMD3\|PSMA4\|PSMA5\|UBA52\|TCEB1\|PSMD14\|PSMB7\|PSMC5\|PSMD12\|PSMD1\|PSME1\|PSME2\|PSMD7 | CP_REACTOME |
| NABA_ECM_GLYCOPROTEINS | 5.84E-05 | 0.01329493 | 0.4451069 | 1.55975693 | 583 | 88 | LAMA4\|FBLN1\|CILP\|OTOG\|SPON2\|FNDC1\|TNC\|LAMB4\|NTN4\|SLIT3\|TINAGL1\|BMPER\|SPARC\|TECTB\|LAMA1\|VWA7\|MFAP5\|SSPO\|FBLN7\|NPNT\|IGFBP7\|NTN5\|TGFBI\|TNN\|SLIT2\|LGI3\|VWDE\|CTGF\|FGL2\|COMP\|THSD4\|SVEP1\|LAMC1\|VWA2\|SRPX\|MATN4\|IGFALS\|LAMC3\|NDNF\|EMILIN2\|VIT\|RSPO3\|GAS6\|FGA\|SRPX2\|USH2A\|HMCN2\|FBLN2\|CRIM1\|HMCN1\|LAMA5\|VWF\|FGB\|IGSF10\|VWCE\|LAMB3\|RSPO2 | CP_CANONICAL_PATHWAYS |
| REACTOME_AUTODEGRADATION_OF_THE_E3_UBIQUITIN_LIGASE_COP1 | 0.00374979 | 0.14651897 | 0.48423077 | 1.54551511 | 37444 | 34 | PSMB10\|PSMB6\|PSMC2\|PSMC4\|PSMB3\|PSMB5\|PSMD8\|PSMB4\|PSMA3\|PSMD3\|PSMA4\|PSMA5\|UBA52\|PSMD14\|PSMB7\|PSMC5\|PSMD12\|PSMD1\|PSME1\|PSME2\|PSMD7 | CP_REACTOME |
| REACTOME_ER_PHAGOSOME_PATHWAY | 0.00354863 | 0.14651897 | 0.47585166 | 1.53433497 | 35454 | 37 | PSMB10\|PSMB6\|PSMC2\|PSMC4\|PSMB3\|PSMB5\|PSMD8\|PSMB4\|PDIA3\|PSMA3\|PSMD3\|PSMA4\|PSMA5\|UBA52\|PSMD14\|PSMB7\|PSMC5\|PSMD12\|PSMD1\|PSME1\|PSME2\|PSMD7 | CP_REACTOME |

| | | | | | | | | |
|---|---|---|---|---|---|---|---|---|
| NABA_CORE_MATRISOME | 1.42E-05 | 0.0093152 | 0.42463893 | 1.52641841 | 141 | 127 | LAMA4\|LUM\|KERA\|COL20A1\|FBLN1\|CILP\|COL22A1\|SPOCK2\|OTOG\|SPON2\|FNDC1\|TNC\|COL4A6\|LAMB4\|NTN4\|SLIT3\|COL4A3\|TINAGL1\|BMPER\|SPOCK3\|SPARC\|TECTB\|LAMA1\|VWA7\|MFAP5\|SSPO\|FBLN7\|OPTC\|NPNT\|IGFBP7\|NTN5\|TGFBI\|TNN\|SLIT2\|LGI3\|VWDE\|DCN\|CTGF\|FGL2\|COMP\|THSD4\|SVEP1\|LAMC1\|VWA2\|COL4A4\|COL4A5\|ASPN\|EPYC\|SRPX\|MATN4\|IGFALS\|LAMC3\|NDNF\|EMILIN2\|VIT\|COL1A2\|RSPO3\|GAS6\|FGA\|SRPX2\|COL8A2\|COL6A3\|USH2A\|HMCN2\|VCAN\|COL13A1\|FBLN2\|CRIM1 | CP_CANONICAL_PATHWAYS |
| REACTOME_P53_INDEPENDENT_G1_S_DNA_DAMAGE_CHECKPOINT | 0.00484155 | 0.14651897 | 0.47526173 | 1.52227664 | 48356 | 35 | PSMB10\|PSMB6\|PSMC2\|PSMC4\|PSMB3\|PSMB5\|PSMD8\|PSMB4\|PSMA3\|PSMD3\|PSMA4\|PSMA5\|UBA52\|PSMD14\|PSMB7\|PSMC5\|PSMD12\|PSMD1\|PSME1\|PSME2\|PSMD7\|PSMD10\|PSMB1\|PSMD6\|CHEK1\|PSMC3 | CP_REACTOME |
| REACTOME_SCF_BETA_TRCP_MEDIATED_DEGRADATION_OF_EMI1 | 0.00496951 | 0.14651897 | 0.47469652 | 1.52046627 | 49634 | 35 | PSMB10\|PSMB6\|PSMC2\|PSMC4\|PSMB3\|PSMB5\|PSMD8\|PSMB4\|PSMA3\|PSMD3\|PSMA4\|PSMA5\|UBA52\|PSMD14\|PSMB7\|PSMC5\|PSMD12\|PSMD1\|PSME1\|PSME2\|PSMD7 | CP_REACTOME |
| REACTOME_SIGNALING_BY_WNT | 0.00355839 | 0.14651897 | 0.46705656 | 1.51968273 | 35563 | 40 | PSMB10\|PSMB6\|PSMC2\|PSMC4\|PSMB3\|PSMB5\|PSMD8\|PSMB4\|PSMA3\|PSMD3\|PSMA4\|PSMA5\|UBA52\|PSMD14\|PSMB7\|PSMC5\|PSMD12\|PSMD1\|CTNNB1\|PSME1\|PSME2\|PSMD7\|PPP2R5D\|PSMD10\|PPP2R5A\|PSMB1 | CP_REACTOME |
| REACTOME_DESTABILIZATION_OF_MRNA_BY_AUF1_HNRNP_D0 | 0.00589813 | 0.14651897 | 0.47076137 | 1.50786185 | 58909 | 35 | PSMB10\|PSMB6\|PSMC2\|PSMC4\|PSMB3\|PSMB5\|PSMD8\|PSMB4\|PSMA3\|PSMD3\|PSMA4\|PSMA5\|UBA52\|PSMD14\|PSMB7\|PSMC5\|PSMD12\|PSMD1\|PSME1\|PSME2\|PSMD7\|PSMD10\|PSMB1\|PSMD6\|PSMC3\|HSPB1 | CP_REACTOME |
| REACTOME_SCFSKP2_MEDIATED_DEGRADATION_OF_P27_P21 | 0.00457037 | 0.14651897 | 0.46460297 | 1.5073412 | 45673 | 39 | PSMB10\|PSMB6\|PSMC2\|PSMC4\|PSMB3\|PSMB5\|PSMD8\|PSMB4\|CKS1B\|CCNA1\|PSMA3\|PSMD3\|PSMA4\|PSMA5\|UBA52\|PSMD14\|PSMB7\|PSMC5\|PSMD12\|PSMD1\|PSME1\|PSME2\|PSMD7\|PSMD10\|PSMB1\|PSMD6\|CDK2\|PSMC3 | CP_REACTOME |
| REACTOME_CROSS_PRESENTATION_OF_SOLUBLE_EXOGENOUS_ANTIGENS_ENDOSOMES | 0.00735008 | 0.147657 | 0.47550899 | 1.50618151 | 73358 | 32 | PSMB10\|PSMB6\|PSMC2\|PSMC4\|PSMB3\|PSMB5\|PSMD8\|PSMB4\|PSMA3\|PSMD3\|PSMA4\|PSMA5\|PSMD14\|PSMB7\|PSMC5\|PSMD12\|PSMD1\|PSME1\|PSME2\|PSMD7 | CP_REACTOME |
| REACTOME_CDT1_ASSOCIATION_WITH_THE_CDC6_ORC_ORIGIN_COMPLEX | 0.00376541 | 0.14651897 | 0.45831818 | 1.50333374 | 37640 | 43 | PSMB10\|PSMB6\|PSMC2\|PSMC4\|PSMB3\|PSMB5\|PSMD8\|PSMB4\|ORC5\|PSMA3\|PSMD3\|PSMA4\|PSMA5\|UBA52\|GMNN\|MCM8\|PSMD14\|PSMB7\|PSMC5\|PSMD12\|PSMD1\|PSME1\|ORC1\|PSME2\|PSMD7 | CP_REACTOME |
| REACTOME_REGULATION_OF_INSULIN_SECRETION | 0.00788346 | 0.147657 | 0.45824356 | 1.47755945 | 78764 | 37 | ADRA2A\|ISL1\|GNG3\|SPCS3\|PRKAR2B\|SPCS1\|GNG8\|SEC11A\|PRKACG\| | CP_REACTOME |

| | | | | | | | | |
|---|---|---|---|---|---|---|---|---|
| | | | | | | | ITPR2\|GNB2\|GATA4\|GNG12\|GNB1\|SLC2A2\|RAPGEF3\|ITPR3\|GNAS\|SLC25A4\|GNG5 | |
| REACTOME_CDK_MEDIATED_PHOSPHORYLATION_AND_REMOVAL_OF_CDC6 | 0.00889195 | 0.147657 | 0.46122765 | 1.47732507 | 88811 | 35 | PSMB10\|PSMB6\|PSMC2\|PSMC4\|PSMB3\|PSMB5\|PSMD8\|PSMB4\|PSMA3\|PSMD3\|PSMA4\|PSMA5\|UBA52\|PSMD14\|PSMB7\|PSMC5\|PSMD12\|PSMD1\|PSME1\|PSME2\|PSMD7\|PSMD10\|PSMB1\|PSMD6\|CDK2\|PSMC3 | CP_REACTOME |
| GO_EXTRACELLULAR_MATRIX | 5.90E-06 | 0.00098235 | 0.39856068 | 1.46588667 | 58 | 191 | LAMA4\|LUM\|KERA\|MMP13\|COL20A1\|FBLN1\|CILP\|COL22A1\|SPOCK2\|GPC2\|SPON2\|TNC\|COL4A6\|PLAT\|TFPI2\|LAMB4\|WNT10B\|NTN4\|SLIT3\|COL4A3\|TINAGL1\|KAZALD1\|SPOCK3\|IL1RL1\|SPARC\|TECTB\|ADAMTS17\|FGFBP3\|LAMA1\|PHOSPHO1\|TGFBR3\|MFAP5\|FBLN7\|OPTC\|NPNT\|IGFBP7\|TGFBI\|TNN\|SLIT2\|CCDC80\|TGFB3\|DCN\|CTGF\|TNFRSF11B\|SFRP2\|DAG1\|COMP\|THSD4\|ALPL\|HSP90B1\|CPA6\|LAMC1\|TIMP4\|APLP1\|WNT11\|ADAMTS1\|VWA2\|WNT3A\|COL4A4\|NDP\|COL4A5\|ASPN\|PCSK6\|EPYC\|TGFB2\|F2\|FGFR2\|MMP20\|CRTAP\|GFOD2\|ADAMTS6\|LECT1\|SERPINE2\|LAMC3\|NDNF\|WNT9A\|FREM3\|CDON\|FREM1\|MMP2\|EMILIN2\|VIT\|ADAMTS18\|COL1A2\|ADAMTSL5\|LOXL2\|ADAMTS16\|COL8A2\|COL6A3\|USH2A\|HMCN2\|VCAN\|GPC5\|ADAMTS10\|MMP24\|CCBE1\|FBLN2 | GO_CELLULAR_COMPONENT |
| REACTOME_AUTODEGRADATION_OF_CDH1_BY_CDH1_APC_C | 0.00860329 | 0.147657 | 0.43704569 | 1.44064927 | 86010 | 45 | PSMB10\|PSMB6\|PSMC2\|PSMC4\|PSMB3\|PSMB5\|PSMD8\|PSMB4\|PSMA3\|PSMD3\|PSMA4\|PSMA5\|UBA52\|UBE2D1\|PSMD14\|PSMB7\|PSMC5\|PSMD12\|ANAPC11\|PSMD1\|PSME1\|PSME2\|ANAPC4\|PSMD7\|PSMD10\|CDC16\|PSMB1\|ANAPC10 | CP_REACTOME |
| REACTOME_CYCLIN_E_ASSOCIATED_EVENTS_DURING_G1_S_TRANSITION_ | 0.00872693 | 0.147657 | 0.43673489 | 1.43962476 | 87246 | 45 | PSMB10\|PSMB6\|PSMC2\|PSMC4\|PSMB3\|PSMB5\|PSMD8\|RB1\|PSMB4\|CKS1B\|CCNA1\|PSMA3\|PSMD3\|PSMA4\|PSMA5\|UBA52\|PSMD14\|PSMB7\|PSMC5\|PSMD12\|PSMD1\|PSME1\|PSME2\|PSMD7 | CP_REACTOME |
| GO_BASOLATERAL_PLASMA_MEMBRANE | 0.0018002 | 0.14986674 | 0.41114809 | 1.4326718 | 18001 | 82 | HPGD\|ENPP1\|LIN7A\|TGFA\|MLC1\|ADRA2A\|KCNQ1\|SLC39A5\|CDH16\|SLC26A5\|LEPR\|CTNNA2\|NUMB\|ATP2B4\|AQP4\|IL6R\|STX4\|HFE2\|PDZD11\|ERBB2\|DAG1\|RHBG\|MEGF11\|ABCC4\|PTH1R\|SLC10A1\|LIN7C\|DLG2\|SLC16A8\|HSP90AB1\|SHROOM4\|SLC2A2\|RAPGEF3\|PKD1\|PKD2\|CAV1\|P2RY12\|CTNNB1\|ABCC1\|CDH17\|DSTYK | GO_CELLULAR_COMPONENT |
| GO_PROTEINACEOUS_EXTRACELLULAR_MATRIX | 6.66E-05 | 0.0073926 | 0.39259174 | 1.43195632 | 665 | 163 | LAMA4\|LUM\|KERA\|MMP13\|FBLN1\|CILP\|COL22A1\|SPOCK2\|GPC2\|SPON2\|TNC\|COL4A6\|TFPI2\|LAMB4\|WNT10B\|NTN4\|SLIT3\|COL4A3\|KAZALD1\|S | GO_CELLULAR_COMPONENT |

| | | | | | | | | |
|---|---|---|---|---|---|---|---|---|
| | | | | | | | POCK3\|IL1RL1\|SPARC\|TECTB\|ADAMTS17\|LAMA1\|PHOSPHO1\|TGFBR3\|MFAP5\|FBLN7\|OPTC\|NPNT\|TGFBI\|TNN\|SLIT2\|CCDC80\|DCN\|CTGF\|TNFRSF11B\|DAG1\|COMP\|THSD4\|ALPL\|CPA6\|LAMC1\|TIMP4\|APLP1\|WNT11\|ADAMTS1\|VWA2\|WNT3A\|COL4A4\|COL4A5\|ASPN\|EPYC\|MMP20\|CRTAP\|GFOD2\|ADAMTS6\|LECT1\|LAMC3\|NDNF\|WNT9A\|FREM3\|FREM1\|MMP2\|EMILIN2\|VIT\|ADAMTS18\|COL1A2\|ADAMTSL5\|LOXL2\|ADAMTS16\|COL8A2\|COL6A3\|USH2A\|HMCN2\|VCAN\|GPC5\|ADAMTS10\|MMP24\|CCBE1\|FBLN2 | |
| REACTOME_SIGNALING_BY_THE_B_CELL_RECEPTOR_BCR | 0.00988422 | 0.147657 | 0.39019399 | 1.35277394 | 98841 | 77 | PSMB10\|PSMB6\|PSMC2\|PIK3CD\|PSMC4\|PSMB3\|PSMB5\|PSMD8\|AKT1S1\|PSMB4\|SH3KBP1\|IKBKG\|PHLPP1\|PIK3AP1\|PSMA3\|PSMD3\|ITPR2\|PSMA4\|PSMA5\|UBA52\|TSC2\|CBLB\|FOXO4\|PSMD14\|PSMB7\|PSMC5\|IKBKB\|ITPR3\|BCL10\|PSMD12\|PSMD1\|PSME1\|PSME2\|PLCG1\|SYK\|PSMD7\|MAPKAP1 | CP_REACTOME |
| REACTOME_HOST_INTERACTIONS_OF_HIV_FACTORS | 0.01027992 | 0.147657 | 0.38767832 | 1.34686923 | 102798 | 79 | ELMO1\|PSMB10\|PSMB6\|PSMC2\|BANF1\|PSMC4\|PSMB3\|PSMB5\|RBX1\|NUP50\|NUP54\|PSMD8\|PSMB4\|PSMA3\|AP2S1\|PSMD3\|NPM1\|PSMA4\|PSMA5\|UBA52\|NUP205\|TCEB1\|PSMD14\|PSMB7\|PSMC5\|KPNA1\|SLC25A4\|HCK\|PSMD12\|PSMD1\|PSME1\|PSME2\|PSMD7\|CCNT1\|PSMD10\|KPNB1\|AP2A1\|NUP88\|PSMB1 | CP_REACTOME |
| GO_EXTRACELLULAR_SPACE | 4.20E-06 | 0.00098235 | 0.35095654 | 1.33308428 | 41 | 429 | CA6\|LUM\|KERA\|IL6ST\|MMP13\|WFDC1\|COL20A1\|APOF\|ENPP1\|FBLN1\|CILP\|CPSF3L\|THBD\|TGFA\|OTOG\|GPC2\|AGRP\|EDN1\|SPON2\|C8B\|FCN3\|IL13RA2\|KLHL17\|TNC\|MSMP\|PLAT\|PCSK2\|RBP4\|TFPI\|QSOX1\|ENO1\|ITIH2\|WNT10B\|FAM132A\|SLIT3\|MASP1\|SEMA3B\|VLDLR\|TINAGL1\|LCP1\|AGR2\|BMPER\|AMN\|BTD\|ENOX1\|USPL1\|TCN2\|S100B\|SPOCK3\|ITIH4\|IL1RL1\|SPARC\|SEMA3D\|PRDX1\|DNAJC9\|KDSR\|LAMA1\|PRDX6\|TGFBR3\|STOM\|CMTM6\|CPN2\|CTSF\|IL6R\|SEMA3G\|BMP3\|ADAM9\|ANGPT4\|SERPINC1\|SSPO\|MTHFD2\|STX4\|TNFRSF1A\|UBB\|IGFBP7\|PODXL\|FIGF\|SMPD1\|HFE2\|HABP2\|TGFBI\|SLIT2\|TGFB3\|CNOT1\|DCN\|CTGF\|LRRC17\|TNFRSF11B\|ATXN10\|TG\|SFRP2\|DAG1\|RALGAPA2\|BTC\|FGL2\|GNL1\|COMP\|CMTM7\|MST1\|ALPL\|CHGA\|EDN2\|PLA2G15\|CLEC11A\|CHRD\|C9\|SST\|CD40\|CPA6\|LAMC1\|GNB2\|TIMP4\|CFD\|TIMM8B\|MDGA1\|BMP15\|SH3BGRL\|PLA2G6\| | GO_CELLULAR_COMPONENT |

| | | | | | | | | |
|---|---|---|---|---|---|---|---|---|
| | | | | | | | WNT11\|ENOX2\|VWA2\|WNT3A\|UBA52\|RNPEP\|NDP\|CLIC1\|IGF2R\|TNFSF13B\|SORD\|SPTBN2\|LCAT\|KRT78\|IDE\|PCSK6\|EPYC\|VEGFC\|SAAL1\|TGFB2\|SULF1\|F2\|MMP20\|IGFALS\|CTSH\|C2\|NCOA5\|PSMC5\|PYY\|FLT1\|CRTAP\|FGF21\|KL\|TPI1\|SERPINE2\|FGF16\|SERPINE3\|MERTK\|WNT9A\|FREM3\|PPT1\|SEMA3C\|HSPD1\|C5\|MMP2\|PLXDC1\|LSR\|COL1A2\|CAT\|METRN\|PRDX4\|FABP3\|OTOP1\|KNG1\|GAS6\|GNL3\|LOXL2\|INHA\|APOA1BP\|SELE\|FGA\|SRPX2\|MFNG\|CFI\|MFI2\|CTSC\|COL6A3\|PCSK9\|NRG3\|NUCB1\|VCAN\|GPC5\|FKTN\|HMOX1\|AGA\|NAPSA\|CETP\|CIB2\|GRIPAP1\|LAMP2\|CCBE1\|PDGFD | |
| REACTOME_PLATELET_ACTIVATION_SIGNALING_AND_AGGREGATION | 0.0098959 | 0.147657 | 0.37800086 | 1.32888792 | 98958 | 92 | VAV3\|DGKE\|ADRA2A\|GNG3\|TLN1\|SPARC\|GP9\|PRKCQ\|STX4\|CALU\|RHOA\|FIGF\|RAC2\|TGFB3\|GNG8\|MGLL\|ITPR2\|GNB2\|CFD\|ABCC4\|RHOB\|GNG12\|VEGFC\|GNB1\|TGFB2\|F2\|RAPGEF3\|ITPR3\|SRC\|VCL\|WDR1\|GNG5\|PIK3CG\|COL1A2\|P2RY12\|KNG1\|PIK3CB\|CDC42\|FGA\|SYK\|HSPA5\|SCG3\|DGKG\|LAMP2 | CP_REACTOME |
| NABA_MATRISOME | 6.08E-05 | 0.01329493 | 0.34646924 | 1.30917435 | 607 | 365 | LAMA4\|LUM\|KERA\|MMP13\|COL20A1\|FBLN1\|CILP\|PLXNA4\|MEGF6\|TGFA\|COL22A1\|SPOCK2\|OTOG\|GPC2\|SPON2\|FCN3\|FNDC1\|TNC\|COL4A6\|PLAT\|PLXNC1\|ITIH2\|LAMB4\|WNT10B\|NTN4\|SLIT3\|COL4A3\|MASP1\|SEMA3B\|TINAGL1\|KAZALD1\|BMPER\|S100B\|SPOCK3\|ITIH4\|SPARC\|TECTB\|SEMA3D\|ADAMTS17\|FGFBP3\|LAMA1\|CPN2\|VWA7\|MFAP5\|CTSF\|P3H1\|TMPRSS15\|SEMA3G\|BMP3\|ADAM9\|MASP2\|ANGPT4\|SERPINC1\|SSPO\|FBLN7\|OPTC\|NPNT\|IGFBP7\|FIGF\|NTN5\|HABP2\|TGFBI\|TNN\|SLIT2\|LGI3\|TGFB3\|VWDE\|BRINP2\|DCN\|CTGF\|FGF13\|P3H2\|SFRP2\|BTC\|FGL2\|S100Z\|FGF14\|COMP\|MST1\|THSD4\|CLEC11A\|CHRD\|CLEC19A\|SVEP1\|MEGF11\|LAMC1\|TIMP4\|BMP15\|WNT11\|ADAMTS1\|VWA2\|WNT3A\|COL4A4\|WIF1\|COL4A5\|P4HA3\|TNFSF13B\|CLEC17A\|FAM20B\|ANGPTL5\|ASPN\|LGALSL\|SCUBE2\|P4HA2\|PCSK6\|EPYC\|PLXNA3\|SRPX\|MATN4\|VEGFC\|OGFOD1\|TGFB2\|SULF1\|F2\|PLOD1\|MMP20\|IGFALS\|ITIH5\|CTSH\|FGF21\|ISM1\|SEMA5A\|ADAMTS6\|SERPINE2\|LAMC3\|FGF16\|SERPINE3\|NDNF\|WNT9A\|FREM3\|SEMA3C\|P4HTM\|FREM1\|MMP2\|EMILIN2\|PLXNB3\|VIT\|PLXDC1\|ADAMTS18\|COL1A2\|SCUBE3\|NTF4\|RSPO3\|A | CP_CANONICAL_PATHWAYS |

| | | | | | | | | |
|---|---|---|---|---|---|---|---|---|
| | | | | | | | DAMTSL5\|KNG1\|GAS6\|LOXL2\|INHA\|FGA\|SRPX2\|ADAMTS16\|COL8A2\|CTSC\|COL6A3\|USH2A\|SDC4\|HMCN2\|NRG3\|VCAN\|GPC5\|ADAMTS10\|MMP24\|COL13A1\|CCBE1\|FBLN2\|PDGFD\|SDC2\|CRIM1\|EGLN3\|CRLF3\|CXCL14\|FGF3 | |
| REACTOME_HEMOSTASIS | 0.001278 | 0.14651897 | 0.35362646 | 1.30603516 | 12779 | 208 | VAV3\|THBD\|DGKE\|P2RX1\|ADRA2A\|PLAT\|TFPI\|MYB\|AKAP1\|KIF3B\|GNG3\|TLN1\|SPARC\|PDE6A\|CDK5\|ITGB2\|KLC3\|GP9\|GRB14\|PRKCQ\|DOCK10\|SLC8A3\|PRKAR2B\|ANGPT4\|SERPINC1\|IRF7\|STX4\|CALU\|RHOA\|FIGF\|PTPN6\|RAC2\|TGFB3\|GNG8\|LRP8\|DOCK8\|PRKACG\|MGLL\|ITPR2\|GNB2\|CFD\|GATA4\|ABCC4\|GUCY1A3\|PRKG1\|ITGAL\|RHOB\|GNG12\|SLC16A8\|VEGFC\|GNB1\|RACGAP1\|TGFB2\|GUCY1B3\|PDE1A\|F2\|RAPGEF3\|PDE5A\|ITPR3\|SRC\|VCL\|WDR1\|GNAS\|GNG5\|MERTK\|PIK3CG\|CABLES1\|AK3\|COL1A2\|CAV1\|P2RY12\|KNG1\|PIK3CB\|PDE3A\|SH2B3\|PLCG1\|CDC42\|PDE6B\|MAFK\|FGA\|SYK\|SH2B1\|KLC2\|HSPA5\|GUCY1A2\|SCG3\|GATA5\|KIFC1\|DOCK5\|ABL1\|DGKG\|SLC7A5\|LAMP2\|MFN2\|PPP2R5D\|GATA6\|PLEK\|SLC7A9\|KIF23\|PRKAR1B\|PPP2R5A | CP_REACTOME |
| REACTOME_TRANSMEMBRANE_TRANSPORT_OF_SMALL_MOLECULES | 0.0056565 | 0.14651897 | 0.34575275 | 1.26923872 | 56564 | 184 | ATP10B\|ABCC9\|SLC9A5\|SLC12A4\|SLC7A2\|ATP8B3\|SLC39A5\|SLC35D2\|SLC15A2\|SLC4A3\|HTR3A\|SLC13A3\|SLC16A7\|GNG3\|ABCA12\|SLC24A1\|NUP50\|SLC27A6\|NUP54\|G6PC3\|AQP4\|SLC8A3\|PRKAR2B\|ATP8A2\|SLC39A4\|SLC31A1\|SLC6A15\|GABRB1\|GNG8\|SLC24A3\|RHAG\|FLVCR1\|SLC6A14\|SLC39A10\|GABRA3\|SLC9A8\|SLC5A2\|SLC2A6\|PRKACG\|SLC38A2\|SLC22A16\|SLC6A3\|SLC17A5\|GNB2\|ABCC4\|SLC39A1\|SLC44A4\|SLC12A1\|NUP205\|HK1\|GNG12\|SLC16A8\|GNB1\|SLC35B4\|SLC2A2\|RAB11A\|ABCC2\|GABRB3\|SLC29A3\|GNAS\|SLCO2B1\|GNG5\|ABCA2\|SLC18A2\|ABCC5\|ATP6V0D1\|SLC5A5\|SLC24A5\|ABCC1\|SLC1A4\|SLC35A2\|ADCY8\|HMOX1\|ATP6V1D\|SLC7A5\|ATP8B4\|ATP11B\|SLC39A2\|CP\|SLC6A2\|SLC35A1\|SLC1A1\|PEX19\|ATP6V0B\|SLC2A10\|SLC7A9\|NUP88\|PRKAR1B\|ABCA7\|SLC26A1\|SLC6A9 | CP_REACTOME |

1
